# Oxylipin carbonyl composition in the chloroplast compartments

**DOI:** 10.1101/2025.11.25.690356

**Authors:** Sergey A. Khorobrykh, Yoko Iijima, Daisuke Shibata, Jun’ichi Mano

## Abstract

Aldehydes and ketones derived from oxidized lipids (oxylipin carbonyls) accumulate in plants under stress conditions, acting as signaling molecules or cytotoxic agents through electrophilic modification of biomolecules. Because chloroplasts generate reactive oxygen species and contain lipids rich in unsaturated fatty acids, they are likely major sites of oxylipin carbonyl formation.

To identify chloroplast-derived carbonyl species and their sub-organellar distribution, we analyzed carbonyls in spinach leaves, whole chloroplasts, thylakoids, envelopes and stroma. A total of 26 carbonyl species were detected, of which 13 were identified. Chloroplasts contained 14 species, including formaldehyde, propionaldehyde, acetaldehyde, acetone, butyraldehyde, (*E*)-2-pentenal, (*Z*)-3-hexenal, (*E*)-2-hexenal, *n*-hexanal and 4-hydroxy-(*E*)-2-hexenal.

Phenylacetaldehyde and *n*-pentanal, which were abundant in leaves, were not associated with chloroplasts. Thylakoid membranes contained all chloroplast carbonyls except acetone, indicating that they are a major site of carbonyl formation. Formaldehyde was highly enriched in thylakoids and envelopes, providing evidence that it can be generated within chloroplasts, most likely from membrane lipids. Estimated concentrations of several carbonyls in membranes reached over 1 millimolar. This high-resolution mapping of oxylipin carbonyls within chloroplast compartments, achieved here for the first time, provides a spatial framework for understanding their formation and functional roles in plant stress responses.

## 1. Introduction

Generation of reactive oxygen species (ROS) such as superoxide radical (O_2•–_) and singlet oxygen (^1^O_2_) is intrinsically associated with redox reactions in photosynthesis and respiration. These ROS are often produced within or in the close vicinity of biomembranes, and as a result, the polyunsaturated fatty acid (PUFA) moieties in a lipid molecule, especially at their pentadienoic structure, are oxidized to form lipid hydroperoxides (LOOHs). LOOHs are subsequently degraded by radical catalysts, such as Fe^2+^ ion or O_2•–_, into various aldehydes and ketones, collectively known as oxylipin carbonyls (Farmer & Mueller 2013). In plants, a wide variety of oxylipin carbonyls occurs, differing in carbon chain length, number and position of unsaturated bonds and degree of oxygenation (Mano 2012, Schauenstein et al. 1977).

Oxylipin carbonyls include molecules with diverse physiological functions. Among them, those containing the *α,β*-unsaturated bond—typically 2-propenal (acrolein), 4-hydroxy-(*E*)-2-nonenal (HNE) and malondialdehyde (MDA)—are collectively termed reactive carbonyl species (RCS) (Mano 2012, Esterbauer et al. 1991), or reactive electrophile oxylipins (Farmer & Mueller 2013). Due to the highly electrophilic nature of the *β*-carbon in these molecules, they are highly cytotoxic. This mechanism is primarily explained by the formation of covalent bonds between RCS and nucleophilic residues of proteins, such as lysine, histidine, and cysteine, as well as guanine bases in nucleic acids (Esterbauer et al. 1991). Recent comprehensive structural analyses of carbonylated plant proteins have revealed that RCS contribute more significantly to protein carbonylation than ROS (Matamoros et al. 2018).

In plants, RCS are major contributors to tissue damage caused by ROS-induced oxidative stress (Mano et al. 2019a, Biswas and Mano 2021). Protein modifications by RCS such as acrolein, HNE, crotonaldehyde, 4-hydroxy-(*E*)-2-hexenal (HHE), and MDA have been reported to increase under salt stress (Mano et al. 2014a). Elevated levels of free RCS have also been observed in leaves under strong light (Mano et al. 2010), seedlings under high salinity (Sultana et al. 2022, Sultana et al. 2024), and roots exposed to toxic concentrations of AlCl₃ (Yin et al. 2010). The direct involvement of RCS in tissue injury is evidenced by the fact that overexpression of RCS-scavenging enzymes or addition of chemical scavengers alleviated stress symptoms in conjunction with suppression of RCS levels (Mano et al. 2005 and 2010, Sultana et al. 2022, Sultana et al. 2024, Yin et al. 2010).

On the other hand, it is becoming increasingly clear that RCS also function as ROS signal transduction molecules in plants. A typical example is the H_2_O_2_-induced programmed cell death (PCD) in tobacco cultured cells (Biswas & Mano 2015). When H₂O₂ was added to trigger PCD, the RCS levels rose before PCD initiation; however, when RCS scavengers were added to suppress this increase, the onset of cell death was inhibited. Meanwhile, the scavengers did not suppress H₂O₂-induced increases in ROS or LOOHs. RCS have also been shown to be involved in the initiation of senescence in siliques and leaves (Nurbekova et al. 2024, Srivastava et al. 2017), and in retrograde signaling from the chloroplast in *Chlamydomonas reinhardtii* (Roach et al. 2017). Furthermore, RCS mediate ROS signals that reinforce auxin signaling for lateral root formation (Biswas et al. 2019), abscisic acid and jasmonic acid signaling for stomata closure (Islam et al. 2016, Islam et al. 2020) and suppression of blue light-induced stomata opening (Murakami et al. 2022).

Accumulating evidence indicates that distinct RCS types exhibit different biological effects, as follows. Acrolein had a stronger effect than HNE on lateral root formation (Biswas et al. 2019), while HNE more effectively inhibited blue light-induced stomatal opening (Murakami et al. 2022). Fumigation of *Arabidopsis thaliana* plants with C₄–C₈ RCS strongly induced heat-responsive genes, whereas acrolein did not (Yamauchi et al. 2015). Thus, RCS exert specific biological effects at low concentrations depending on their type, while exhibiting toxicity at high intracellular concentrations. In vitro toxicity to photosynthesis was strongest for acrolein, then HNE followed, while crotonaldehyde, the RCS that is only one-carbon longer than acrolein, was much less toxic than these two RCS (Mano et al. 2009). Thus, RCS exhibit toxicity at high intracellular concentrations, while exerting specific biological effects at low concentrations depending on their type. For normal physiological activity in plant cells, it is essential to precisely regulate the levels of different oxylipin carbonyls and RCS, which are determined by their flux of production and elimination specifically. Many carbonyl-scavenging and RCS-scavenging enzymes have been identified, and their physiological importance has been elucidated (Mano et al. 2019a, Mano et al. 2019b). However, the mechanisms underlying RCS generation in plant cells remain largely unclear.

To clarify the mechanism of oxylipin carbonyl formation, we focused in this study on identifying the sites of their generation in the chloroplast. There are several reasons to focus on chloroplasts: (1) Chloroplasts are major sources of ROS in leaf tissues. Singlet oxygen (¹O₂) is continuously produced at PSII (Vass et al. 1992, Macpherson et al. 1993), and superoxide radical (O₂•⁻) at PSI (Asada & Kiso 1973). (2) Most lipids in mesophyll cells are found in chloroplasts, and chloroplast lipids are rich in PUFAs. For example, in spinach leaves, PUFAs account for 85% of total fatty acids in thylakoids (Garab et al. 2017). Mène-Saffrané et al. (2007) observed that the *A. thaliana vte* mutant, which lacks *α*-tocopherol, accumulated more MDA (measured as thiobarbituric acid-reactive substances) than the wild type. Thus, even under physiological light intensities, oxidation of thylakoid lipids and generation of oxylipin carbonyls can proceed if ROS are not efficiently scavenged. (3) Chloroplast proteins are potential targets of RCS (Mano et al. 2014a, Mano et al. 2009, Ji et al. 2023). (4) LOOHs may be generated on the donor side of PSII when the water-splitting complex is inactivated (Khorobrykh et al. 2011). (5) Thylakoid and envelope membranes contain several lipoxygenases (LOXs) that catalyze lipid peroxidation, and hydroperoxide lyase (HPL), which decomposes LOOHs into oxylipin carbonyls, is also localized in the envelope membrane (Blée and Joyard 1996).

Lipid peroxidation induced by ROS proceeds predominantly via two distinct mechanisms. The first entails the electrophilic addition of ^1^O_2_ to a double bond within an unsaturated fatty acid, yielding an endoperoxide intermediate that is subsequently converted to a LOOH. The second involves the abstraction of a hydrogen atom from a PUFA by hydroxyl or alkoxyl radicals, initiating a free radical chain reaction that results in the formation of multiple LOOH molecules. Nevertheless, the extent to which these pathways contribute to LOOH generation *in vivo* remains uncertain, as current knowledge is largely derived from studies on the thermal oxidation of edible oils (Grosch 1987).

LOOHs are also formed via the catalysis by LOXs. LOXs are thought to be latent; they are activated only on certain physiological occasions such as disruption of cells due to mechanical wounding of a tissue (Mochizuki et al. 2016). An activated LOX catalyzes the addition of an O_2_ molecule to a specific position(s) of a PUFA molecule. Plants have multiple isoforms of LOX (for example, *A. thaliana* has six), but most of them have not been investigated for their enzymatic properties and thus it is largely unclear which isozymes contribute, and to what extents they do, to the formation of LOOHs under varying environmental conditions.

Resulting LOOHs are cleaved by some redox catalysts into carbonyl radicals, which then abstract H atom from a neighboring molecule (likely a lipid molecule in the membrane) to be a carbonyl molecule (i.e. an aldehyde or a ketone). HPL can cleave specific types of LOOH to form specific types of oxylipin carbonyls.

We previously analyzed the composition of oxylipin carbonyls in the leaves of *A. thaliana fad7fad8* mutant, which lacks trienoic fatty acid biosynthesis in the chloroplast, and compared it with the wild type (Col-0) (Mano et al. 2014b). The mutant had significantly lower levels of RCS such as acrolein, crotonaldehyde and (*E*)-2-pentenal, as well as non-RCS carbonyls such as formaldehyde and acetaldehyde, than the wild type. In contrast, carbonyls such as acetone and *n*-hexanal were found at higher levels in the mutant. This suggests that the oxylipin carbonyls contained less in the mutant were derived from the trienoic fatty acids in chloroplasts, i.e. linolenic acid and 16:3 fatty acid. However, the relevance of the two most abundant aldehydes, formaldehyde and acetaldehyde, to lipid oxidation remains uncertain, as they have generally been considered byproducts of alcohol metabolism. Furthermore, since carbonyls were extracted from whole leaves in the above-mentioned study, it is still unknown whether specific carbonyl species were produced inside or outside the chloroplast. The distinction between thylakoid- and envelope-derived carbonyls—i.e., which species originate from which membranes—also has yet to be clarified.

In this study, we analyzed oxylipin carbonyls in chloroplast subcompartments, including the whole chloroplast, thylakoids, envelopes, and stroma. We found distinct differences in carbonyl composition among these compartments and estimated the local concentrations of specific carbonyls within each. To our knowledge, this represents the first high-resolution analysis of oxylipin-derived carbonyls at the sub-chloroplast level.

## 2. Methods

### 2.1. Chemicals

Acetonitrile, formic acid and tetrahydrofuran used for carbonyl analysis were of HPLC grade. 2,4-Dinitrophenylhydrazine (DNPH) was recrystallized from its ethanol solution. Other reagents were of analytical grade. DNPH-II Eluent B for carbonyl analysis was obtained from Fujifilm Wako Pure Chemical (Osaka, Japan).

### 2.2. Isolation of chloroplasts, and preparation of thylakoid, envelopes and the stroma fraction

Intact chloroplasts were isolated from field-grown *Spinacia oleracea* L. leaves. All procedures were done under ice-chilled temperature. Thirty grams of leaves without midrib were ground 3 times for 5 s each with a blender in 250 mL of isolation medium containing 330 mM sorbitol, 50 mM Hepes-KOH, pH 7.6, 2 mM diethylenetriaminepentaacetic acid, 1 mM MnCl_2_, 1 mM MgCl_2_. Bovine serum albumin (0.5% (w/v)) and ascorbic acid (5 mM) were added to the isolation medium just before homogenization. The homogenate was filtered through two layers of Miracloth (Merck Millipore, Tokyo, Japan) and centrifuged at 1,100 × *g* for 5 min. The pellet was resuspended in the isolation medium, and carefully loaded onto Percoll (Cytiva, Tokyo, Japan) density gradient (top to bottom: 7 mL of 40% Percoll, 3 mL of 80% Percoll) made in the isolation medium. After centrifugation at 1,630 × *g* for 15 min using a swing rotor, intact chloroplasts forming a band at the 40%–80% interface of Percoll were collected and washed twice with a medium containing 330 mM sorbitol, 50 mM Hepes-KOH, pH 7.8, 4 mM MgCl_2_ and centrifuged for 5 min at 1,100 × *g*. This preparation should contain the outer and inner envelopes, stroma, and thylakoids and hereafter we designate it ’whole chloroplast’.

The envelopes and thylakoids were isolated from the whole chloroplast preparation (1.5 mg Chl) by a gentle osmotic shock in 1.5 mL hypotonic medium containing 0.05 mM sucrose, 10 mM Hepes-KOH, pH 7.8, 4 mM MgCl_2_ and purified by sucrose gradient centrifugation according to Douce and Joyard (1982). Envelope membranes, accumulated at the interface between 0.6 M and 0.93 M sucrose, and thylakoids (including both the thylakoid membranes and the lumen) at the 0.93 M/1.5 M interface were collected and suspended in the hypotonic medium to final volume of 1.5 mL. The upper layer on top of the 0.6 M sucrose layer (1.5 mL) was collected as the stroma fraction.

### 2.3 Carbonyl analysis

From 1 g of spinach leaves that were removed of midrib, carbonyls were extracted with 10 mL of acetonitrile (HPLC grade) containing 12.5 nmol of 2-ethylhexanal as the internal standard (I. S.) and 0.005% (w/v) butylhydroxytoluene at 60°C for 30 min. Carbonyls from the whole chloroplasts, thylakoids, envelopes and the stroma fraction were extracted by adding 1.5 mL suspension of a sample (see Methods 2.2.) to 10 mL acetonitrile (HPLC grade) containing 12.5 nmol of 2-ethylhexanal as the internal standard (I. S.) and 0.005% (w/v) butylhydroxytoluene. After an incubation at 60°C for 30 min, the mixture was centrifuged at 1500 × *g* for 10 min using the R4S/39 swing rotor (Eppendorf Himac Technologies Co., Ltd., Hitachinaka, Japan) and the supernatant was collected as the extract. Carbonyls were extracted in acetonitrile at 60°C incubation in the absence of added acid or base. where substantial release of protein-bound carbonyls is not expected. Thus, the measured carbonyls predominantly represent free species. The extract (about 10 mL) was mixed with 0.2 mL of 25 mM DNPH in acetonitrile and 0.191 mL of formic acid (99%), and incubated for 60 min at 25°C to derivatize carbonyls to the hydrazones, which were then purified through a Bond Elut C18 cartridge (Agilent Technologies Japan, Hachioji) as previously reported (Matsui et al. 2009). A blank sample, containing no plant material but the I. S., was prepared through the same procedure used for plant material. The obtained DNP-derivatives of carbonyls were separated by a reverse-phase HPLC with the Wakosil DNPH-II column (Fujifilm Wako) and detected by absorbance at 340 nm (Matsui et al. 2009) and by Fourier transform ion cyclotron mass spectrometry using Finnigan LTQ-FR (Thermo Electron, Rockford, Il, USA) (Mano et al. 2014b). Carbonyls were identified based on their retention time (RT) as compared with the derivatives of authentic carbonyls (Matsui et al. 2009). Identification of carbonyls was also confirmed with MS/MS analysis as previously described (Mano et al. 2014b). DNP-carbonyl peaks of which the RT did not match any of authentic standards were designated ’unidentified carbonyls’. We used Xcalibur 2.2 software (Thermo Fisher Scientific, Tokyo, Japan) to assign elemental compositions to DNP–carbonyl derivatives for each peak based on high-resolution MS data of the deprotonated molecular ion [M – H]⁻. The molecular formula of the corresponding carbonyl was calculated from that of the DNP–carbonyl derivative by reversing the DNPH condensation reaction (addition of H₂O and subtraction of DNPH, C₆H₆N₄O_4_).

The amount of each carbonyl was calculated from its peak area detected at 340 nm. This wavelength choice is a practical compromise to detect various DNP-carbonyls with different absorption spectra with a single wavelength monitor (Matsui et al. 2009). Briefly, the area of each carbonyl peak (*A*_s_) was normalized to the area of the I.S. peak (*A*_IS_) (i.e., *A*_s_/*A*_IS_ was obtained). The peaks in the blank sample (*A*_b_) were also normalized to its *A*_IS_ and then net peak area of a carbonyl normalized to *A*_IS_ was obtained (*A*_s_/*A*_IS_ – *A*_b_/ *A*_IS_). The amount of an identified carbonyl in nmol was calculated according equation (*A*_s_/*A*_IS_ – *A*_b_/ *A*_IS_) × 12.5 nmol × *k*, where 12.5 nmol is the amount of I. S. added to the sample and *k* is a conversion factor, which is specific to distinct known carbonyl species (Matsui et al. 2009). For unidentified carbonyls the conversion factor was taken as 1. Some peaks of unidentified carbonyls, when examined with LC/MS, consisted of two signals of different *m*/*z* values. The relative contents of distinct carbonyls in such a peak were calculated based on the intensity ratio of their molecular ion signals, assuming that the ionization efficiency of the two DNP-carbonyls were the same.

## 3. Results

### 3.1. Carbonyls in spinach leaves

In living cells, oxylipin carbonyls are present either in free forms or in protein-bound adducts (Esterbauer et al. 1991). Protein-bound forms, including both Michael adducts and Schiff base adducts, are stable or irreversible due to subsequent rearrangement or cross-linking (Abuja & Esterbauer 1995, Waeg et al. 1996, Doorn & Peterson 2003, Del Rio et al. 2005).

Hence, the carbonyl content determined in this study predominantly represents free carbonyls and the contribution of protein-bound form is expected to be small. Indeed, when both free carbonyl extractable with acetonitrile and carbonyl-modified proteins were determined in salt-stressed plants, their levels did not correlate each other (Mano et al. 2014).

Typical chromatograms of DNP-derived carbonyls from a leaf sample and a blank are shown in Fig. 1. A large peak eluted around 2.3 min in the blank chromatogram (broken line) was of DNPH. A substantial amount of unreacted free DNPH remained also in the plant sample. Several DNP-derived carbonyls were eluted before and immediately after the free DNPH peak; however, this region was not analyzed due to poor resolution between peaks. When a small carbonyl peak appeared as a shoulder on another larger peak, the slope of the larger peak was taken as the baseline for quantification of the minor peak. For example, in Fig. 1, the HHE peak (RT 7.09 min) appeared as a shoulder of the larger acetaldehyde peak (RT 6.59 min). In this case, the area of the HHE peak was measured using the slope of acetaldehyde peak as the baseline. Similarly, the acetone peak (RT 8.97 min) and (*E*)-2-pentenal peak (RT 18.36 min) were treated as shoulders on lager peaks.

**Figure 1.**
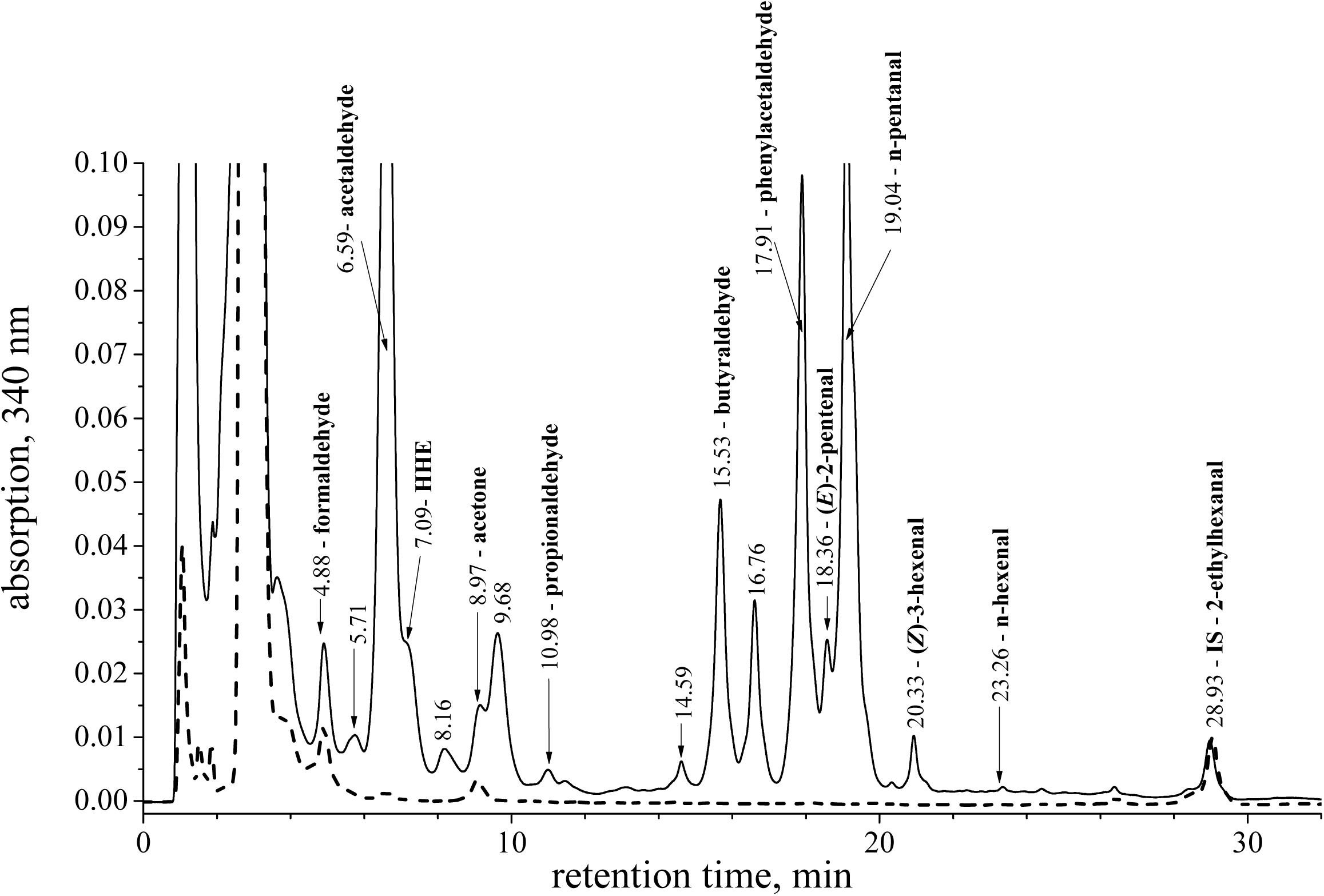
A typical chromatogram of DNP-derivatized carbonyls in field-grown spinach leaves. The dashed line represents the chromatogram of a blank sample.

#### 3.1.1 Identified carbonyls

In total, 26 carbonyls were detected across leaves, chloroplasts, and chloroplast subcompartments (Tables 1 and 2), among which 13 species were identified based on their retention times and MS/MS spectra (Supplementary Figs. S1 & S2). Table 1 summarized the leaf carbonyls from two samples harvested on two different dates. All detected carbonyls normalized on fresh below 100 nmol (g FW)^−1^. Several carbonyls were abundant in both samples, i.e., acetaldehyde (71.6 nmol (g FW)^−1^), formaldehyde (44.5 nmol (g FW)^−1^) and acetone (18.3 nmol (g FW)^−1^). Some carbonyls showed relatively high mean values, but substantial variation between sampling dates. These include (*E*)-2-hexenal (39.2 nmol (g FW)^−1^), *n*-pentanal (36.1 nmol (g FW)^−1^), phenylacetaldehyde (Phe-al) (21.6 nmol (g FW)^−1^), HHE (17.0 nmol (g FW)^−1^), and butyraldehyde (11.6 nmol (g FW)^−1^). Other carbonyls, such as propionaldehyde, (*E*)-2-pentenal, (*Z*)-3-hexenal, *n*-hexanal and *n*-nonanal were present at relatively low levels.

**Table 1.** Carbonyls detected in spinach leaves. Carbonyl contents are expressed as nmol (g FW of leaves)⁻¹. Values represent mean ± SE. Day 1 (*n* = 4) and Day 2 (*n* = 3) indicate independent biological experiments. Overall means were estimated using a linear mixed-effects model (LMM). n.d., not detected. * Trace level (peak detected by MS analysis, but not quantifiable). The *m/z* value of the molecular ion [M – H]^−^ represents the corresponding DNP-derived carbonyl. Blank samples (without plant material) contained formaldehyde, acetaldehyde, and acetone at 1.86 ± 0.21, 0.05 ± 0.002, and 1.3 ± 0.14 nmol mL^−1^, respectively.

| RT<br>(min) | *m/z* | Carbonyl species (Formula)<sup>a</sup> | Day 1 | Day 2 | Overall mean |
| --- | --- | --- | --- | --- | --- |
| nmol carbonyl (g FW)<sup>-1</sup> |  |  |  |  |  |
| **Identified** |  |  |  |  |  |
| 4.88 | 209.03 | Formaldehyde (CH<sub>2</sub>O) | 31.07 ± 1.65 | 58.33 ± 9.57 | 44.5 ± 13.6 |
| 6.59 | 223.05 | Acetaldehyde (C<sub>2</sub>H<sub>4</sub>O) | 81.29 ± 10.28 | 60.68 ± 7.31 | 71.6 ± 10.3 |
| 7.09 | 293.09 | HHE (C<sub>6</sub>H<sub>10</sub>O<sub>2</sub>) | 7.36 ± 2.69 | 26.92 ± 6.67 | 17.0 ± 9.7 |
| 8.97 | 237.06 | Acetone (C<sub>3</sub>H<sub>6</sub>O) | 25.47 ± 4.31 | 10.8 ± 0.78 | 18.3 ± 7.3 |
| 10.98 | 237.06 | Propionaldehyde (C<sub>3</sub>H<sub>6</sub>O) | 1.75 ± 0.25 | 1.06 ± 0.07 | 1.41 ± 0.35 |
| 15.53 | 251.08 | Butyraldehyde (C<sub>4</sub>H<sub>8</sub>O) | 20.25 ± 1.21 | 2.91 ± 0.72 | 11.6 ± 8.6 |
| 17.91 | 299.08 | Phenylacetaldehyde (C<sub>8</sub>H<sub>8</sub>O) | 41.35 ± 3.06 | 1.88 ± 0.22 | 21.6 ± 19.7 |
| 18.36 | 263.08 | (*E*)-2-Pentenal (C<sub>5</sub>H<sub>8</sub>O) | 9.12 ± 0.91 | 2.56 ± 0.33 | 5.85 ± 3.38 |
| 19.04 | 265.09 | *n*-Pentanal (C<sub>5</sub>H<sub>10</sub>O) | 70.1 ± 5.49 | 2.08 ± 0.36 | 36.1 ± 34 |
| 20.31 | 277.09 | (*Z*)-3-Hexenal (C<sub>6</sub>H<sub>10</sub>O) | 1.59 ± 0.27 | 8.77 ± 2.45 | 5.14 ± 3.6 |
| 22.07 | 277.09 | (*E*)-2-Hexenal (C<sub>6</sub>H<sub>10</sub>O) | n.d. | 78.43 ± 7.92 | 39.2 ± 39.2 |
| 23.26 | 279.11 | *n*-Hexanal (C<sub>6</sub>H<sub>12</sub>O) | 0.19 ± 0.03 | 0.8 ± 0.02 | 0.49 ± 0.3 |
| 35.15 | 321.16 | *n*-Nonenal (C<sub>9</sub>H<sub>18</sub>O) | 1.04 ± 0.2 | 1.43 ± 0.13 | 1.22 ± 0.19 |
| **Unidentified** |  |  |  |  |  |
| 5.71 | 281.05 | #1 (C<sub>4</sub>H<sub>6</sub>O<sub>3</sub>) | 1.15 ± 0.41 | 2.18 ± 0.15 | 1.65 ± 0.52 |
| 7.09 | 295.10 | #2 (C<sub>6</sub>H<sub>12</sub>O<sub>2</sub>) | n.d. (trace) | 26.4 ± 2.1 | 13.2 ± 13.2 |
| 8.16 | 377.15 | #3 (C<sub>11</sub>H<sub>18</sub>O<sub>3</sub>) | 6.71 ± 2.45 | 20.06 ± 1.29 | 13.3 ± 6.67 |
| 8.16 | 291.07 | #4 (C<sub>6</sub>H<sub>8</sub>O<sub>2</sub>)<sup>b</sup> | n.d. | n.d. | n.d. |
| 8.16 | 295.10 | #5 (C<sub>6</sub>H<sub>12</sub>O<sub>2</sub>) | n.d. | n.d. * | n.d. |
| 9.68 | 315.07 | #6 (C<sub>8</sub>H<sub>8</sub>O<sub>2</sub>) | 24.6 ± 2.28 | 0.43 ± 0.06 | 12.5 ± 12.1 |
| 9.68 | 301.06 | #7 (C<sub>7</sub>H<sub>6</sub>O<sub>2</sub>) | 2.98 ± 0.55 | 0.5 ± 0.06 | 1.76 ± 1.24 |
| 10.15 | 309.08 | #8 (C<sub>6</sub>H<sub>10</sub>O<sub>3</sub>) | n.d. | n.d. * | n.d. |
| 14.59 | 403.16 | #9 (C<sub>13</sub>H<sub>20</sub>O<sub>3</sub>) | 2.34 ± 0.65 | 0.7 ± 0.06 | 1.54 ± 0.82 |
| 15.98 | 391.16 | #10 (C<sub>12</sub>H<sub>20</sub>O<sub>3</sub>) | n.d. | 0.66 ± 0.32 | 0.32 ± 0.32 |
| 16.76 | 338.09 | #11 (C<sub>10</sub>H<sub>9</sub>ON) | 13.42 ± 1.47 | 0.51 ± 0.07 | 6.98 ± 6.95 |
| 16.76 | 403.16 | #12 (C<sub>13</sub>H<sub>20</sub>O<sub>3</sub>) | 6.53 ± 0.31 | 12.47 ± 0.1 | 9.5 ± 2.97 |
| 21.08 | 335.17 | #13 (C<sub>10</sub>H<sub>20</sub>O) | n.d. | 9.26 ± 1.55 | 4.6 ± 4.6 |

<sup>a</sup> Formula of an unidentified carbonyl was deduced from the *m/z* value of the DNP-derivative.
<sup>b</sup> Unidentified carbonyl #4 was detected only in the thylakoids (Table 2), but is listed here for showing its RT and *m/z* value.

**Table 2.** Distribution of carbonyls in sub-chloroplast compartments. Whole chloroplasts isolated from spinach leaves were fractionated into thylakoids (including the lumen), envelopes, and stroma. Carbonyl contents in each compartment are expressed as amounts per unit chloroplast content (nmol (mg Chl)⁻¹) of the original chloroplast preparation to allow direct comparison among compartments. Chloroplasts used for compartment analysis were prepared from leaves harvested on two independent days (*n* = 2 per day). Mean values and standard errors (SE) were estimated using a linear mixed-effects model (LMM). n.d., not detected by absorbance. Asterisks indicate that the mass signal was detected, but the peak area was not quantifiable due to very low absorbance (trace level).

| Carbonyl species | Carbonyl content per unit chloroplast contents (nmol (mg Chl)<sup>-1</sup>) |  |  |  |
| --- | --- | --- | --- | --- |
|  | Whole chloroplast | Thylakoids | Envelopes | Stroma |
| Formaldehyde | 36.3 ± 13.3 | 12.65 ± 2.93 | 2.12 ± 0.68 | 2.88 ± 1.78 |
| Acetaldehyde | 1.77 ± 0.83 | 2.50 ± 1.11 | n.d. * | 1.29 ± 0.3 |
| HHE | 0.13 ± 0.02 | 0.98 ± 0.17 | 0.42 ± 0.16 | 1.47 ± 0.17 |
| Acetone | 0.84 ± 0.44 | n.d. * | 0.64 ± 0.4 | 0.68 ± 1.3 |
| Propionaldehyde | 0.20 ± 0.03 | 0.41 ± 0.09 | n.d. | 0.17 ± 0.04 |
| Butyraldehyde | 0.41 ± 0.23 | 0.35 ± 0.19 | n.d. | 0.66 ± 0.54 |
| Phenylacetaldehyde | n.d. * | n.d. * | n.d. | n.d. |
| (*E*)-2-Pentenal | 0.69 ± 0.27 | 0.67 ± 0.11 | 0.18 ± 0.02 | 0.45 ± 0.09 |
| *n*-Pentanal | n.d. (trace) | 0.09 ± 0.02 | n.d. | n.d. |
| (*Z*)-3-Hexenal | 5.30 ± 1.3 | 24.9 ± 7.7 | 7.47 ± 0.9 | 35.2 ± 6.5 |
| (*E*)-2-Hexenal | 4.30 ± 1.56 | 15.04 ± 10.5 | 0.63 ± 0.2 | 17.2 ± 3.6 |
| *n*-Hexanal | 0.18 ± 0.04 | 0.20 ± 0.12 | 0.18 ± 0.1 | 0.95 ± 0.36 |
| **Unidentified** |  |  |  |  |
| #1 (C<sub>4</sub>H<sub>6</sub>O<sub>3</sub>) | 0.50 ± 0.21 | 0.33 ± 0.16 | n.d. * | n.d. * |
| #2 (C<sub>6</sub>H<sub>12</sub>O<sub>2</sub>) | n.d. | n.d. | n.d. | n.d. |
| #3 (C<sub>11</sub>H<sub>18</sub>O<sub>3</sub>) | 11.88 ± 0.32 | 8.61 ± 2.0 | n.d. | n.d. |
| #4 (C<sub>6</sub>H<sub>8</sub>O<sub>2</sub>) | n.d. | 0.4 ± 0.18 | n.d. * | n.d. |
| #5 (C<sub>6</sub>H<sub>12</sub>O<sub>2</sub>) | n.d. * | 0.16 ± 0.08 | n.d. | n.d. * |
| #6 (C<sub>8</sub>H<sub>8</sub>O<sub>2</sub>) | n.d. | n.d. | n.d. | n.d. |
| #7 (C<sub>7</sub>H<sub>6</sub>O<sub>2</sub>) | n.d. * | n.d. | n.d. | n.d. |
| #8 (C<sub>6</sub>H<sub>10</sub>O<sub>3</sub>) | n.d. * | 1.05 ± 0.44 | n.d. * | n.d. |
| #9 (C<sub>13</sub>H<sub>20</sub>O<sub>3</sub>) | n.d. * | n.d. * | n.d. | n.d. * |
| #10 (C<sub>12</sub>H<sub>20</sub>O<sub>3</sub>) | 0.29 ± 0.09 | 0.29 ± 0.05 | n.d. * | n.d. * |
| #11 (C<sub>10</sub>H<sub>9</sub>ON) | n.d. | n.d. | n.d. | n.d. |
| #12 (C<sub>13</sub>H<sub>20</sub>O<sub>3</sub>) | 1.08 ± 0.10 | 0.75 ± 0.12 | n.d. | n.d. |
| #13 (C<sub>10</sub>H<sub>20</sub>O) | n.d. | n.d. | n.d. | n.d. |

Most of these carbonyls have been also detected in the leaves of *A. thaliana* and tobacco (Mano et al. 2012). One characteristic feature of spinach leaves is the relatively high level of Phe-al. This aldehyde is formed from phenylalanaine (Phe), and the high Phe content in spinach compared with other vegetable plants (Nemzer et al. 2021) may account for its elevated level. In contrast, several RCS such as acrolein, MDA, crotonaldehyde, methacrolein, (*E*)-2-hexenal and (*E*)-2-heptenal, which had been detected in *A. thaliana* and tobacco, were present only at very low or undetectable levels in spinach.

#### 3.1.2. Unidentified carbonyls

In addition to identified carbonyls, several DNP-derivatized carbonyl peaks did not match any available standards. MS analysis indicated that some peaks contained more than one carbonyl species, and a total of 13 unidentified carbonyls were distinguished, numbered #1-#13 according to their elution order. Their molecular formulas and the retention times are listed in Table 1. The levels of these unidentified carbonyls, estimated under the assumption that the molar absorption coefficient of their DNP-derivatives was equivalent to that of DNP-I. S., were comparable to those of identified carbonyls. Of the 13 unidentified carbonyls, twelve were detected in leaves. Some of them showed large variation between sampling dates. For example, #2 (C_6_H_12_O_2_) was the most abundant carbonyl in Day 2 sample, but was not detected in Day 1 sample. The #13 carbonyl (C_10_H_20_O) showed similar tendency. In contrast, unidentified #6 and #11 were more contained in the Day 1 sample.

The MS/MS spectra of the unidentified carbonyls are shown in Supplementary Fig. S3. Unidentified carbonyl #1 is a ketone, as its DNP-derivative did not produce the *m*/*z* 163 fragment, which is characteristic of aldehydes under these conditions (Mano et al. 2014b). Candidate compounds for C_4_H_6_O_3_ include 3-oxobutanoic acid or an α-keto acid, such as keto-pyruvic acid. Unidentified carbonyls #6 (C₈H₈O₂) may correspond to 4-hydroxyphenylacetaldehyde, which can be produced via a tyrosine-derived route (Xu et al. 2020), whereas #11 (C₁₀H₉ON) may correspond to indole-3-acetaldehyde, a tryptophan-derived intermediate in the auxin biosynthesis in plants (Mano and Nemoto 2012). For the unidentified carbonyls #8 (C_12_H_20_O₃) and C_11_H_18_O_3_ (*m/z* 377.14), candidate compounds generated via lipid oxidation are 12-oxo-(*Z*)-9-dodecenoic acid and 11-oxo-(*Z*)-9-undecenoic acid, respectively, which are formed via the ^1^O_2_-dependent oxidation of (*Z*)*-*10-hexadecenoic acid (Montillet et al. 2004). These results provide a basis for comparison with chloroplast carbonyl profiles described below.

### 3.2. Distinction of carbonyls of chloroplast- and non-chloroplast origins

To distinguish between carbonyl species derived from chloroplasts and those derived from other cellular sources, chloroplasts were isolated from spinach leaves (designated as the “whole chloroplast” preparation), and their carbonyl contents were determined. Figure 2 shows a representative chromatogram of DNP-derivatized carbonyls in the whole chloroplast. A total of 20 carbonyl species were detected, 12 of which were identified (Table 2). Some of these compounds were detected only at trace levels by MS analysis. Major carbonyls included formaldehyde (∼36 nmol (mg Chl)^−1^), (*Z*)-3-hexenal (∼5 nmol (mg Chl) ^−1^), (*E*)-2-hexenal (∼4 nmol (mg Chl) ^−1^) and acetaldehyde (∼2 nmol (mg Chl)^−1^).

**Figure 2.**
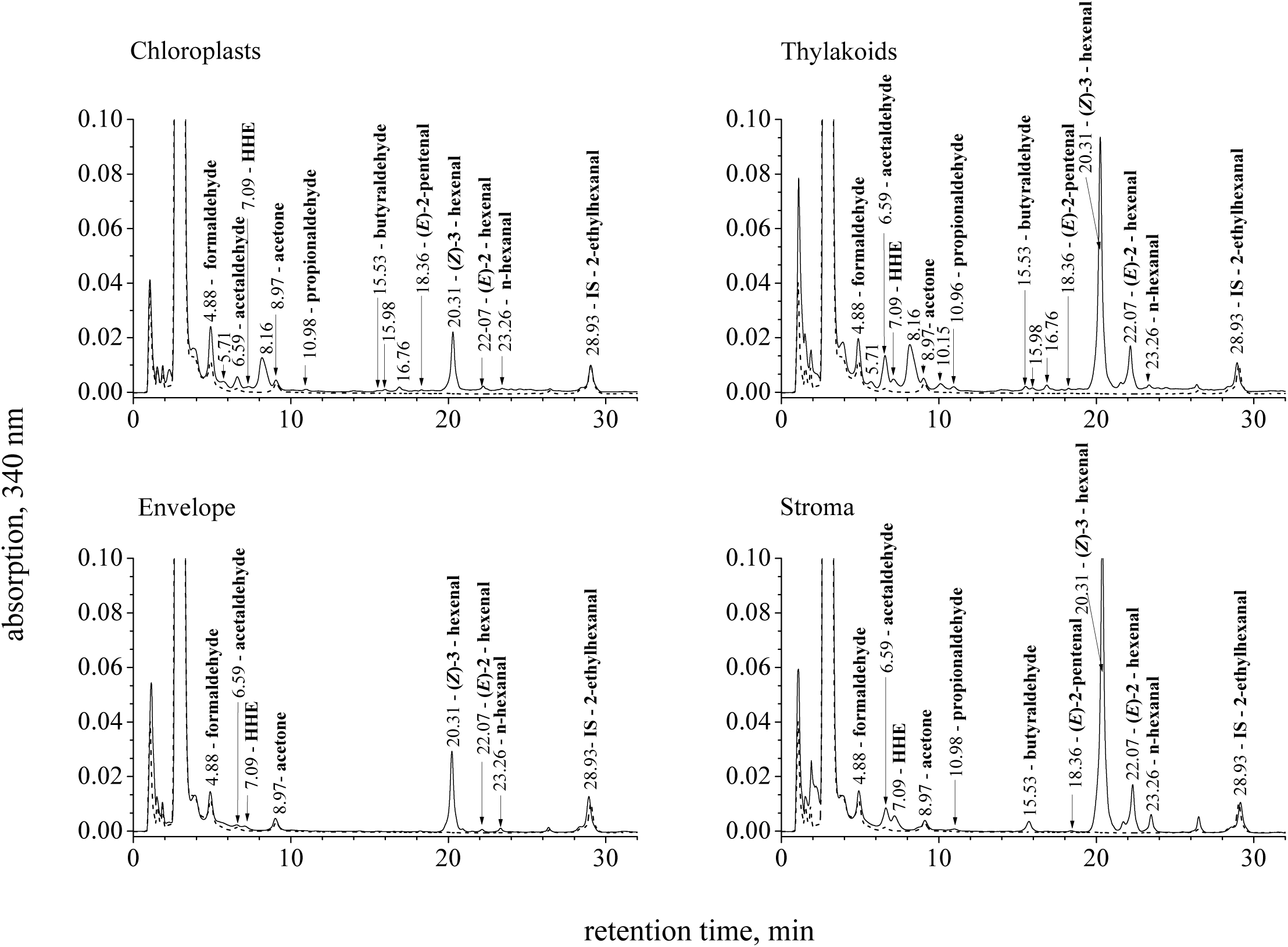
Typical chromatograms of DNP-derivatized carbonyls in chloroplasts, thylakoid membranes, the chloroplast envelope, and the stroma of chloroplasts. The dashed line represents the chromatogram of a blank sample.

The carbonyl profile in the whole chloroplast differed markedly from that of leaves. Phe-al, which was abundant in leaves, was detected in whole chloroplasts only at trace levels by MS analysis, clearly indicating that it does not originate from chloroplasts. In plants, Phe-al is generated enzymatically in the cytosol via the decarboxylation and oxidation of Phe, which is synthesized in the chloroplast and exported to the cytosol (Kaminaga et al. 2006, Widhalm et al. 2015). The absence of Phe-al in the chloroplast is consistent with this pathway. Similarly, *n*-pentanal, another major aldehyde in leaves, was detected only at trace levels in the chloroplast, suggesting that its primary origin lies outside the chloroplast. Among the unidentified carbonyls, #2 (C_6_H_12_O_2_), #6 (C_8_H_8_O_2_), #7 (C_7_H_6_O_2_), #9 (C_13_H_20_O_3_), #11 (C_10_H_9_ON) and #13 (C_10_H_20_O) were absent or present only at trace levels in chloroplasts and their subcompartments, suggesting non-chloroplast origins. In comparison with leaf profiles (Table 1), the whole chloroplast contained relatively lower levels of acetaldehyde, acetone, propionaldehyde, butyraldehyde, (*E*)-2-pentenal, and HHE. These carbonyls may still be produced within the chloroplast, as the whole chloroplast preparations were essentially free from cytosolic contamination, as indicated by the absence of Phe-al. Therefore, these carbonyls can also be produced within the chloroplast.

Taken together, these observations indicate that a subset of carbonyl species is likely generated within chloroplasts. These include formaldehyde, unidentified #3 (C_11_H_18_O_3_), (*Z*)-3-hexenal, (*E*)-2-hexenal, acetaldehyde, unidentified #12 (C_13_H_20_O_3_), acetone, butyraldehyde, unidentified #1 (C_4_H_6_O_3_), (*E*)-2-pentenal, unidentified #10 (C_12_H_20_O_3_), *n*-hexanal and HHE.

### 3.3. Carbonyls in different chloroplast compartments

To elucidate the origin of chloroplast-associated carbonyls, we analyzed their contents in the thylakoids (including the lumen), envelopes and the stroma. Intact chloroplasts were osmotically ruptured and fractionated into sub-chloroplast compartments by density-gradient centrifugation (see Materials and Methods for details). Representative chromatograms of DNP-derivatized carbonyls in these fractions are shown in Fig. 2.

These fractions were obtained from the same chloroplast preparations used for the analysis of the whole chloroplast. To compare carbonyl contents among compartments, we normalized the detected amounts of carbonyls to the unit amount of original chloroplast preparations. We employed chlorophyll amount (mg Chl) to represent of the amount of chloroplast preparations (Table 2). Table 2 summarizes the mean value and standard error obtained from chloroplasts isolated on two independent dates (*n* = 2 per date).

Thylakoids contained nearly all carbonyl species detected in the whole chloroplast, with the exception of acetone. Notably, thylakoids contained the highest formaldehyde amount in the chloroplast, suggesting that the thylakoid membrane is a major site of its accumulation within the chloroplast. Several unidentified carbonyls (#1, #3, #10 and #12) were detected predominantly or exclusively in the thylakoids. In addition, a new unidentified carbonyl #4 (C_6_H_8_O_2_) was detected only in the thylakoid fraction at the peak with RT 8.16, overlapping the unidentified carbonyls #3 and #5.

In the envelopes, only seven types of carbonyls were found at significant levels. These were dominated by acetone and C_6_-carbonyls, including (*Z*)-3-hexenal, (*E*)-2-hexenal, *n*-hexanal, and HHE (Table 2). The carbonyl profile of the envelopes differs clearly from that of thylakoids. First, acetone was present at relatively high levels in the envelopes while it is absent in thylakoids. Second, acetaldehyde, propionaldehyde and butyraldehyde were contained only at trace levels in the envelopes.

The stroma contained ten types of identified carbonyls at significant levels. Similar to the envelopes, the stromal carbonyl profile was characterized by the presence of C_6_-carbonyls, i.e. (*Z*)-3-hexenal (*E*)-2-hexenal and HHE.

The sum of carbonyl contents in the three compartments—thylakoids, envelops and stroma—would be expected to correspond to the total carbonyl content in the whole chloroplast. However, for several carbonyl species, the combined amounts in the three compartments exceeded those measured in the whole chloroplast. Typical examples include the C_6_-aldehyde (*Z*)-3-hexenal, (*E*)-2-hexenal, HHE and unidentified #5 (C_6_H_12_O_2_), whose contents in the thylakoids were several-fold higher than those in the whole chloroplast. This discrepancy suggests that these carbonyls were generated additionally during the fractionation process, as a result of lipid oxidation by LOXs, which are activated by membrane disruption (Matsui 2006). Conversely, for certain carbonyls such as formaldehyde, the sum of the three compartments was lower than the content measured in the whole chloroplast. This discrepancy may reflect partial loss of these carbonyl species during compartment preparation, possibly due to volatilization or redox reactions.

The unidentified carbonyls #1 (C_4_H_6_O_3_), #3 (C_11_H_18_O_3_), #4 (C_6_H_8_O_2_), #8 (C_12_H_20_O_3_) and #10 (C_13_H_20_O_3_), were distributed solely to the thylakoids. Their contents were not changed during compartment isolation.

### 3.4. Estimation of the carbonyl concentrations in distinct chloroplast compartments

To gain further insight into the physiological significance of oxylipin carbonyls in chloroplasts, we estimated the local concentrations of carbonyls in each compartment. The carbonyl contents in each compartment (nmol (mg Chl)^−1^) obtained in Table 2 were divided by the specific volumes of the corresponding compartments (L (mg Chl)^-1^), which were obtained based on literature values, as follows. The volumes of a single chloroplast have been reported to range between 31 × 10^-15^ L and 44 × 10^-15^ L (Zellnig et al. 2004) and a mean value of 37.5 ×10^-15^ L was used in this study. Given that approximately 10^9^ chloroplasts correspond to 1 mg chlorophyll (Pfanz & Heber 1986), the total chloroplast volume per mg chlorophyll was estimated to be 3.75 ×10^-5^ L (mg Chl)^-1^. Based on reported fractional volumes of chloroplast compartments (Zellnig et al. 2004, Tolleter et al. 2024), the specific volumes were estimated as follows: 8.5×10^−6^ L (mg Chl)^-1^ for thylakoids (including lumen), 1.46 ×10^-6^ L (mg Chl)^-1^ for the envelopes and 2.28 ×10^-5^ L (mg Chl)^-1^ for the stroma.

The estimated local concentrations are summarized in Table 3. These values were calculated from the data in Table 2 and the compartment volumes described above. Because some carbonyls may have been generated or lost during chloroplast fractionation, these estimates should be interpreted with caution and considered to represent the potential capacity of each compartment to accumulate these carbonyls rather than their exact *in vivo* concentrations.

**Table 3.** Estimation of carbonyl concentrations in the chloroplast, the thylakoid membrane, the envelopes and the stroma. Concentration (M) of each carbonyl in distinct compartments was calculated from its content (nmol per mg Chl; Table 2) on the specific volumes (L per mg Chl) obtained from literature (see Results). n.d., Not detected.

| Carbonyl species | Carbonyl concentration (μM) in |  |  |  |
| --- | --- | --- | --- | --- |
|  | Whole chloroplast | Thylakoids | Envelopes | Stroma |
| Formaldehyde | 830 | 1300 | 1200 | 110 |
| Acetaldehyde | 40 | 250 | n.d. | 48 |
| Acetone | 19 | n.d. | 370 | 25 |
| Propionaldehyde | 4.5 | 40 | n.d. | 6.3 |
| Butyraldehyde | 9.3 | 35 | n.d. | 25 |
| (*E*)-2-Pentenal | 16 | 68 | 110 | 170 |
| *n*-Pentanal | n.d. | 9.1 | n.d. | n.d. |
| *n*-Hexanal | 4.1 | 20 | 110 | 0.35 |
| (*Z*)-3-Hexenal | 120 | 2500 | 4400 | 1300 |
| (*E*)-2-Hexenal | 98 | 1500 | 370 | 640 |
| HHE | 2.9 | 99 | 250 | 55 |
| **Unidentified** |  |  |  |  |
| #1 (C<sub>4</sub>H<sub>6</sub>O<sub>3</sub>) | 11 | 33 | n.d. | n.d. |
| #3 (C<sub>11</sub>H<sub>18</sub>O<sub>3</sub>) | 270 | 870 | n.d. | n.d. |
| #4 (C<sub>6</sub>H<sub>8</sub>O<sub>2</sub>) | n.d. | 40 | n.d. | n.d. |
| #5 (C<sub>6</sub>H<sub>12</sub>O<sub>2</sub>) | n.d. | 16 | n.d. | n.d. |
| #8 (C<sub>6</sub>H<sub>10</sub>O<sub>3</sub>) | n.d. | 110 | n.d. | n.d. |
| #10 (C<sub>12</sub>H<sub>20</sub>O<sub>3</sub>) | 6.6 | 29 | n.d. | n.d. |
| #12 (C<sub>13</sub>H<sub>20</sub>O<sub>3</sub>) | 25 | 76 | n.d. | n.d. |

In the whole chloroplast, the most abundant carbonyls were formaldehyde (∼0.83 mM), (*Z*)-3-hexenal (∼0.12 mM), (*E*)-2-hexenal (∼0.1 mM) and the unidentified carbonyl #3 (C_11_H_18_O_3_) (∼0.3 mM). In the thylakoids, carbonyl concentrations varied from approximately 10 µM to 2.5 mM, with (*Z*)-3-hexenal (∼2.5 mM), (*E*)-2-hexenal (∼1.5 mM), formaldehyde (∼1.3 mM) and the unidentified #2 (C_11_H_18_O_3_) (∼1 mM) being the most abundant. In the envelopes, the carbonyls with the highest concentrations were formaldehyde (∼1.2 mM) and (*Z*)-3-hexenal (∼4.4 mM), followed by acetone (∼0.4 mM), (*E*)-2-hexenal (∼0.4 mM), HHE (∼0.25 mM), *n*-hexanal (∼0.1 mM) and (*E*)-2-pentenal (0.1 mM). Notably, acetone accumulated to relatively high levels in the envelopes compared with low abundance in other compartments. In the stroma, (*Z*)-3-hexenal (∼1.3 mM) and (*E*)-2-hexenal (∼0.64 mM) were detected at relatively high concentrations, whereas other carbonyls were present at lower than observed for thylakoids and envelopes.

## 4. Discussion

### 4.1. Overview of major findings

The importance of oxylipin carbonyls in plant oxidative signaling has become increasingly evident in recent years (Biswas & Mano 2021, Knieper et al. 2023). However, mechanistic understanding of their physiological roles remains limited, largely due to a lack of information on their sites of formation and local concentrations. To address this limitation, we conducted a comprehensive analysis of carbonyl species in spinach leaves and chloroplast subcompartments, leading to several key findings. First, the carbonyl composition of chloroplasts was clearly distinct from that of whole leaves (Tables 1 & 2). Several major carbonyls detected in leaves, such as phenylacetaldehyde and acetaldehyde, were absent or present only at trace levels in chloroplasts, indicating that they originate primarily from non-chloroplast sources.

Second, within chloroplasts, carbonyl species exhibited distinct distributions among subcompartments. Many carbonyls were enriched in thylakoids, whereas envelopes and the stroma contained more limited subsets, indicating compartment-specific formation and/or accumulation (Table 2). Notably, formaldehyde was highly enriched in thylakoids and envelopes. Formaldehyde is one of the most abundant carbonyls in plant tissues and is increased in response to various stressors (Mano et al. 2010, Biswas & Mano 2015, Mano et al. 2014b, Sultana et al. 2022), but its intracellular source has been unraveled. Our current results demonstrate, for the first time, that this hydrophilic aldehyde is generated in the chloroplast from membrane lipids.

Third, an estimation of local concentrations of carbonyl species within chloroplast compartments is provided. Previous analyzes of carbonyls in plants have generally assumed a uniform distribution within tissues and yielded sub-millimolar values even under stress conditions (Yin et al. 2010; Sultana et al. 2022). In the present study, millimolar levels in thylakoids and envelopes were estimated for formaldehyde and several C_6_-aldehydes (Table 3). These estimates do not represent their in vivo concentrations because the concentrations of C_6_ carbonyls include values affected by artifacts arising from chloroplast disruption. Nevertheless, the data indicate that certain carbonyl species can reach millimolar range in membranes within chloroplasts. This finding provides a quantitative basis for future investigations into the physiological roles of carbonyl species and the sensitivity of their molecular targets in stressed leaves.

### 4.2. Formation mechanisms of oxylipin carbonyls in the chloroplast

In the following discussion, we consider the mechanisms responsible for the formation of several selected carbonyls. As described above, 17 types of carbonyls including seven unidentified were found in the chloroplast. These chloroplast-associated carbonyls are grouped into two. The first group includes C_5_- and C_6_-carbonyls, including unidentified carbonyls #4 (C_6_H_8_O_2_), #5 (C_6_H_12_O_2_) and #8 (C_6_H_10_O_3_), of which formation mechanisms from lipid peroxides are relatively well-documented. The second group are shorter (C_1_–C_4_) carbonyls, which have been much less considered as lipid-derived. This group includes formaldehyde, acetaldehyde, propionaldehyde, butyraldehyde and acetone. We will not discuss unidentified carbonyls #1 (C_4_H_6_O_3_), #3 (C_11_H_18_O_3_), #10 (C_12_H_20_O_3_) and #12 (C_13_H_20_O_3_) because their structures are not sure. Alternative mechanisms of carbonyl formation that do not involve lipid origins, such as oxidation of alcohols catalyzed by dehydrogenases (Jardine and McDowell 2023), are not considered here.

#### 4.2.1. C_5_- and C_6_-carbonyls

The contents of (*Z*)-3-hexenal and (*E*)-2-hexenal in thylakoids were greater than that in the whole chloroplast by approximately 3-5 times (Table 2), suggesting additional formation of these C_6_ aldehydes during the preparation of sub-chloroplast compartments. These increases are likely attributable to the wound-induced stimulation of lipid peroxidation via lipoxygenases (LOXs). LOXs, which catalyze the addition of O_2_ to a specific site in an unsaturated fatty acid, generating a lipid hydroperoxide (LOOH), are localized to thylakoid membranes and envelopes (Blée & Joyard 1996, Farmaki et al. 2006). LOXs are latent enzymes and activated when Ca^2+^ concentration is raised under mechanical disruption of membranes (Mochizuki & Matsui 2018) or oxidative stress (Thakur & Udayashankar 2019). The resulting LOOHs then derive a series of C_6_ aldehydes such as (*Z*)-3-hexenal, (*E*)-2-hexenal and HHE and the C_5_-alkenal (*E*)-2-pentenal, as illustrated in Fig. 3. LOX-catalyzed oxygenation of linolenic acid forms 13-hydroperoxo-octadecatrienoic acid (13-HPOTE). The bond between C-12 and C-13 of 13-HPOTE can be cleaved non-enzymatically to form (*Z*)-3-hexenal radical, which is then reduced by neighboring molecules to (*Z*)-3-hexenal (Grosch 1987) (Fig. 3). HPL localized in the envelopes (Blée & Joyard 1996) can catalyze the (*Z*)-3-hexenal formation from 13-HPOTE (Matsui 2006).

**Figure 3.**
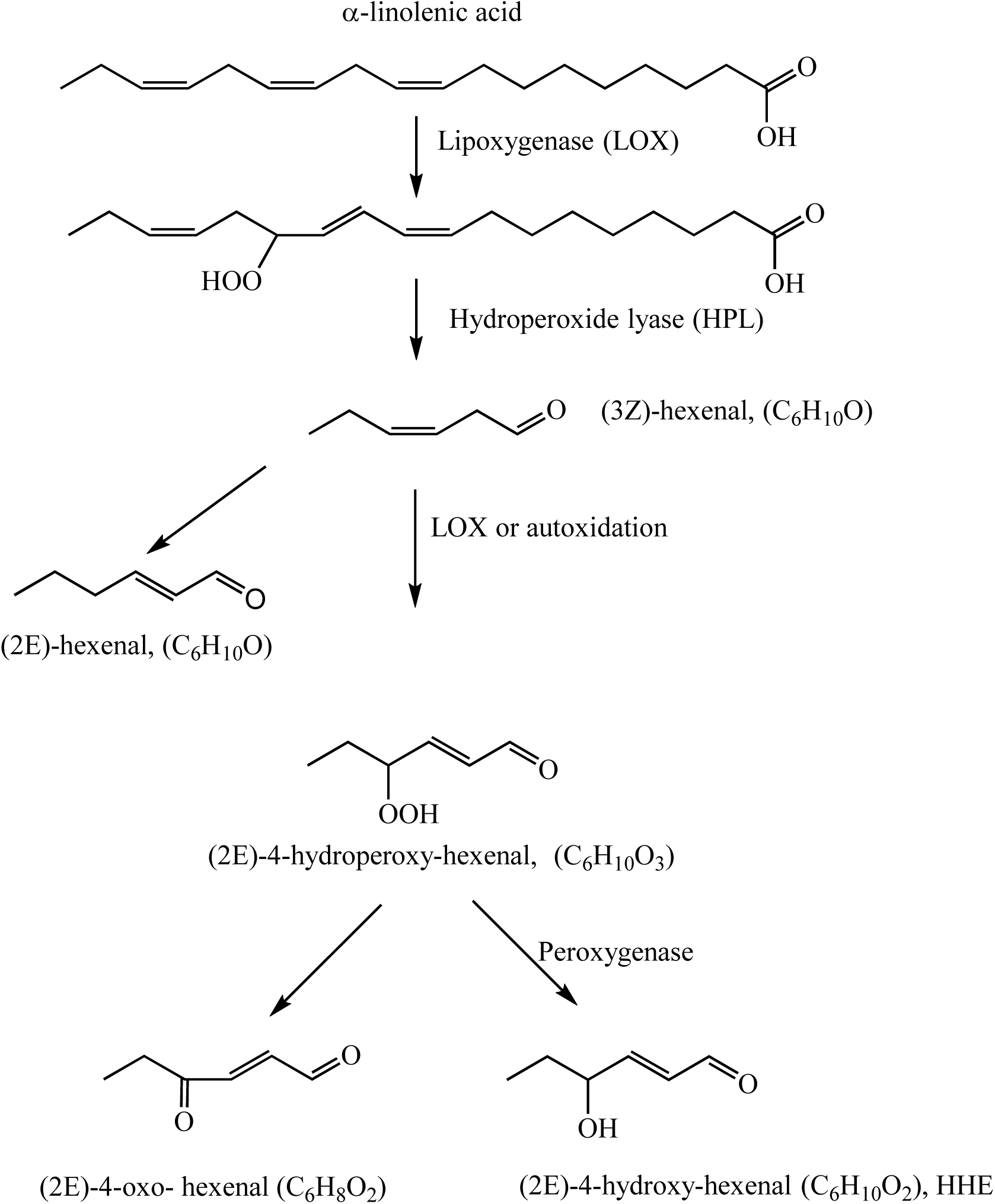
Proposed carbonyls species forming in chloroplasts and chloroplast’s compartments result from oxidation of α-linolenic acid via LOX pathway.

Resulting (*Z*)-3-hexenal can be isomerized to (*E*)-2-hexenal, or oxygenated to 4-hydroperoxy-(*E*)-2-hexenal (C_6_H_8_O_2_), from which HHE and 4-oxo-(*E*)-2-hexenal (C_6_H_10_O_3_) are derived (Matsui 2006). The unidentified carbonyls #4 (C_6_H_8_O_2_) and #8 (C_6_H_10_O_3_) probably are derivative compounds corresponding to the latter two hexenals (Guillén & Goicoechea 2008). 13-HPOTE can be also cleaved at the bond between C-13 and C-14 non-enzymatically, eventually forming (*E*)-2-pentenal (Tawfik et al. 2017). Although not illustrated here, a similar scheme starting with linoleic acid (18:2) through 13- hydroperoxo-octadecadienoic acid will generate *n*-hexanal. High levels of C_6_-carbonyls in the stroma suggest that these carbonyls are released from thylakoid membranes continuously.

#### 4.2.2. C_1_- to C_4_-carbonyls

##### 4.2.2.1 Formaldehyde

Possible mechanisms of its formation from LOOHs have been proposed based on heat-induced oxidation of unsaturated fatty acids such as oleic, linoleic and linolenic acids, in which formaldehyde arises as a secondary degradation product of various LOOHs (Schaich 2013).

Further studies are warranted to determine whether this mechanism operates in chloroplast membrane lipids. While the generation of formaldehyde from LOOHs has been reported in *in vitro* model lipid systems, this provides evidence that formaldehyde can be derived from lipids *in vivo*.

##### 4.2.2.2 Acetaldehyde, propionaldehyde and butyraldehyde

Within chloroplasts, acetaldehyde, propionaldehyde and butyraldehyde were detected in both the thylakoids and the stroma (Table 2). Because they were undetected in the envelopes, these aldehydes are primarily generated in the thylakoids.

Fig. 4 illustrates possible pathways to acetaldehyde and propionaldehyde from linolenic acid. The addition O_2_ to linolenic acid at the C_16_ position will form 16-HPOTE, either via direct adduction of ^1^O_2_ or via hydroxyl radical-initiated chain reactions (Montillet et al. 2004, Cao et al. 2014). Reduction of this peroxide produces the corresponding alkoxyl radical, which undergoes α-scission to form acetaldehyde or propionaldehyde.

**Figure 4.**
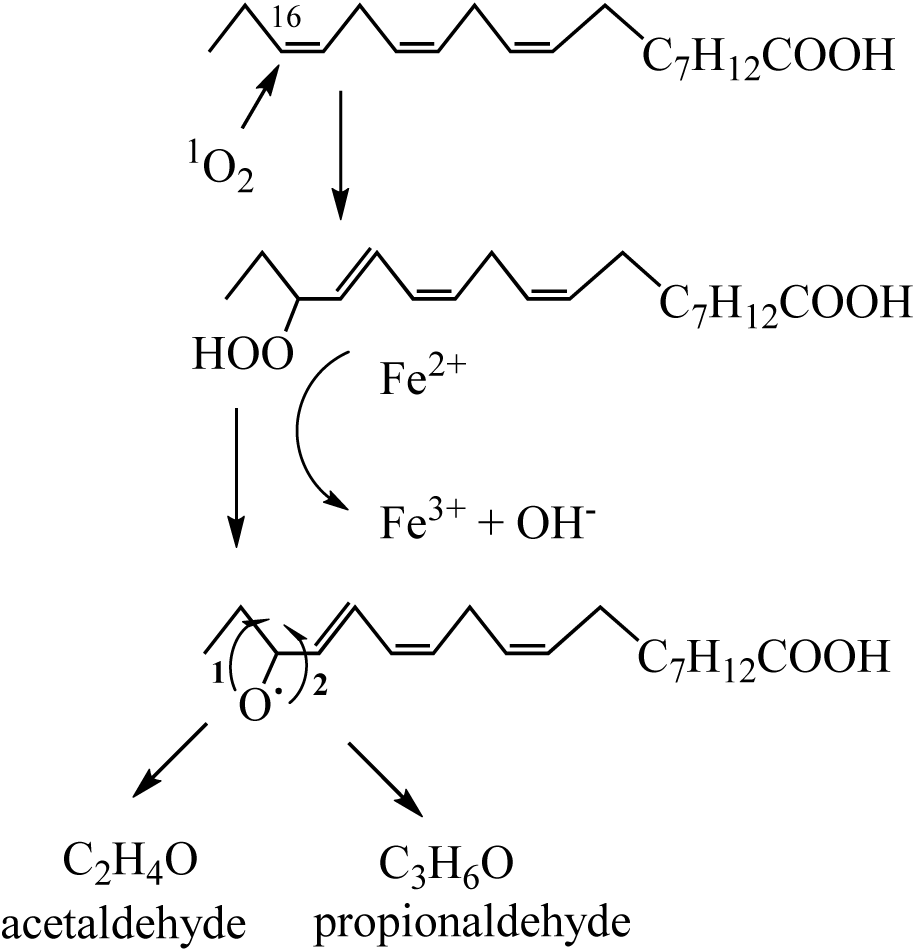
Proposed routes of β-scission of fatty acid hydroperoxides formed via oxidation of α-linolenic acid by singlet oxygen. In route 1, non-allylic C–C bond, carbon-carbon bond that are not located adjacent to a carbon-carbon double bond (C=C), would be cleaved to form acetaldehyde. In route 2, the allylic C–C bond, a single covalent bond between two carbon atoms where one of the carbons is directly bonded to the carbon atoms of a carbon-carbon double bond (C=C), would be cleaved to form propionaldehyde.

The butyraldehyde production mechanism in plants has not been reported. Considering that it was contained more in the *fad7fad8* mutant than in Col-0 line of *A. thaliana*, this aldehyde is preferentially derived from linoleic acid (Mano et al. 2014b). It can be also generated as a secondary degradation product of 13-HPODE (Schaich 2013).

##### 4.2.2.3 Acetone

Acetone was concentrated in the envelopes to a relatively high concentration 370 µM. Its presence in the stroma at 25 µM was likely from the envelopes. A mechanism of acetone formation from unsaturated fatty acids has been proposed based on *in vitro* studies of margarine oxidation, in which acetone is generated through the cleavages of an endoperoxide (Fruehwirth et al. 2021). It is yet to be clarified whether this mechanism also works in the envelope membranes. Interestingly, in contrast, thylakoids did not contain this ketone at detectable levels (Table 2). This implies the presence of a mechanism that suppress endoperoxide accumulation in this compartment. Alternatively, acetone formation may occur via an as-yet-unknown biochemical mechanism specific to the envelope membranes.

### 4.3. Potential physiological roles of chloroplast carbonyls

Plants lacking chloroplast-localized carbonyl-scavenging enzymes exhibit increased susceptibility to photodamage (Yamauchi et al. 2012), implying significant impacts of oxylipin carbonyls on the functions of this organelle. As outlined in the Introduction, several chloroplast proteins have been identified as targets of RCS. Thylakoid membrane proteins such as LHCII and PsbP are modified by MDA (Yamauchi et al. 2008; Kumar et al. 2023), while cyclophilin 20-3 is modified by 4-hydroxy-(*E*)-2-nonenal (HNE) (Mano et al. 2014). In addition, thiol-regulated enzymes of the Calvin cycle, e.g., phosphoribulokinase and glyceraldehyde-3-phosphate dehydrogenase, are inhibited by various carbonyl species (Mano et al. 2009). Among these, acrolein and HNE are known to exhibit particularly high cytotoxicity. More recently, it has been reported that HNE modifies the nascent D1 protein and thereby interferes with its incorporation into the PSII complex (Ji et al. 2023), further emphasizing the vulnerability of photosynthetic machinery to carbonyl stress.

In the present study, we found that formaldehyde accumulates in thylakoid membranes and envelopes at millimolar levels. Compared with highly reactive RCS such as acrolein and HNE, formaldehyde exhibits relatively low cytotoxicity (Reynolds 1977) and a weaker inhibitory effect on photosynthesis (Mano et al. 2009). However, given its substantially higher abundance in chloroplasts, its overall impact on the function of the above-mentioned molecular targets should not be overlooked. Investigation of the effects of formaldehyde on chloroplast components will be an important subject.

Another carbonyl species of particular interest is HHE. This linolenic acid-derived aldehyde has reactivity comparable to that of HNE, which is derived from linoleic acid and is well known for its toxicity. As HHE is present at higher levels than HNE in chloroplasts, it may exert effects on chloroplast components that are comparable to or even greater than those of HNE. The levels of both formaldehyde and HHE increase in plants are significantly increased under environmental stress (Sultana et al. 2022). Evaluation of their effects on photosynthesis will provide important insights into early events of the leaf damage process under stress conditions.

In addition to their cytotoxic effects, oxylipin carbonyls may function as retrograde signals originating from chloroplasts (Roach et al. 2017). In the *A. thaliana* crumpled leaf (*crl*) mutant, which exhibits enhanced singlet oxygen (¹O₂) production due to defects in chloroplast division, localized cell death in leaves requires the presence of C_16_ unsaturated fatty acids in the chloroplast lipids. This observation suggests that oxylipin carbonyls derived from these fatty acids may act as retrograde signals that initiate cell death (Li et al. 2020). Identifying which specific carbonyl species generated in thylakoids serve as the principal retrograde signals will be an important subject for future investigation.

Given these diverse potential roles, the concentrations of individual oxylipin carbonyl species are likely to be tightly regulated within the chloroplast. However, the mechanisms responsible for their detoxification and homeostasis in this plastid remain largely unresolved. Identification of the enzymes involved and characterization of their substrate specificities will be essential for elucidating how the physiological functions of oxylipin carbonyls are controlled in leaves.

## Software

Software used for data processing and preparation of figures are Origin Pro 2015, Thermo Xcalibur version 2.2, and ChemDraw version 19.0.

## CRediT authorship contribution statement

Sergey Khorobrykh: Writing – original draft, Writing – review& editing, Methodology, Data analysis, Conceptualization. Yoko Iijima: LC/MS analysis, Data curation. Daisuke Shibata:

Writing – review& editing. Jun’ichi Mano: Writing – review& editing, Methodology, Supervision, Project administration, Funding acquisition, Data analysis, Conceptualization.

## Declaration of generative AI and AI-assisted technologies in the manuscript preparation process

During the preparation of this work the authors used ChatGPT plus in order to improve the English language of the manuscript. After using this tool, the authors carefully reviewed and edited the text as necessary and take full responsibility for the final content of the publication.

## Data availability

Data will be made available on request.

## Acknowledgements

Stay of S.A.K. in Japan was supported by Goho International Life Science Fund and by Grant-in-Aid of the Japan Society for the Promotion of Science (KAKENHI Grant No. 24H00504).

## Deposition of a preprint

This manuscript has been deposited as a preprint at bioRxiv (DOI: 10.1101/2025.11.25.690356).

---

Supplementary Figure S1. MS/MS spectra of the identified carbonyls.

Supplementary Figure S2. MS/MS spectra of the standard carbonyls.

Supplementary Figure S3. MS/MS spectra of the unidentified carbonyls.

