## Supplementary Fig. S1 for "Oxylipin carbonyl composition in the chloroplast compartments"

RT (min) - 4.88, $[M-H]^-$ = 209.03

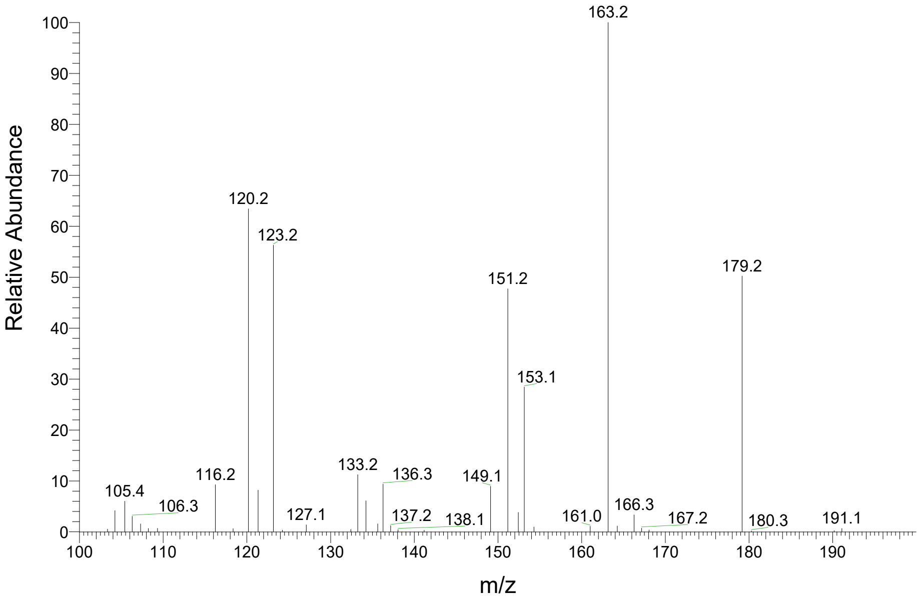


RT (min) - 6.59, $[M-H]^-$ = 223.05

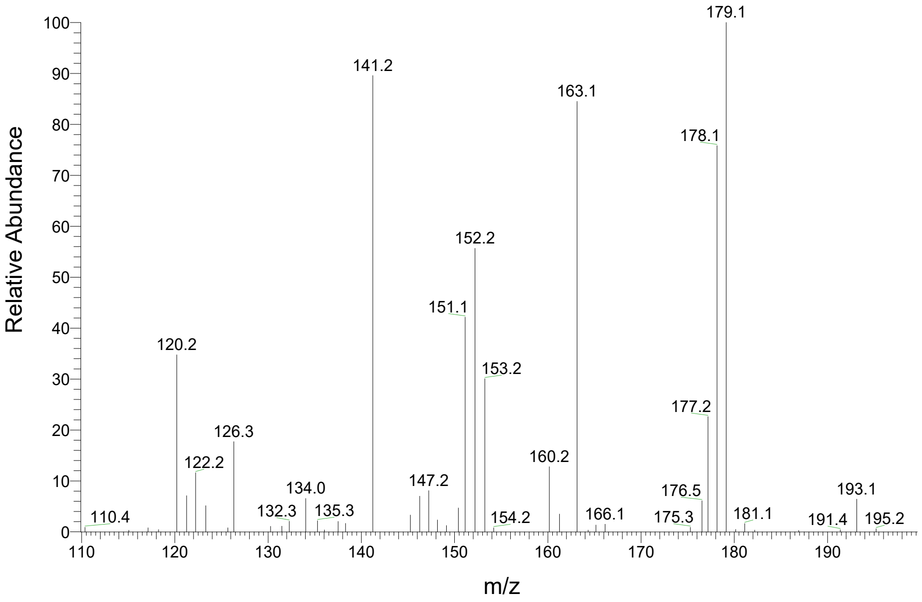


RT (min) - 7.09, $[M-H]^-$ = 293.09

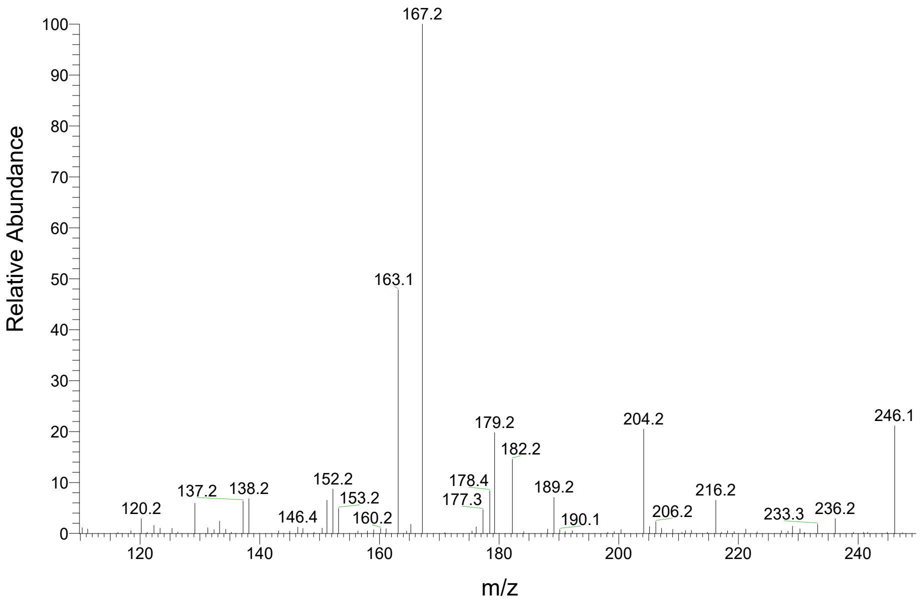


RT (min) - 8.97, $[M-H]^-$ = 237.06

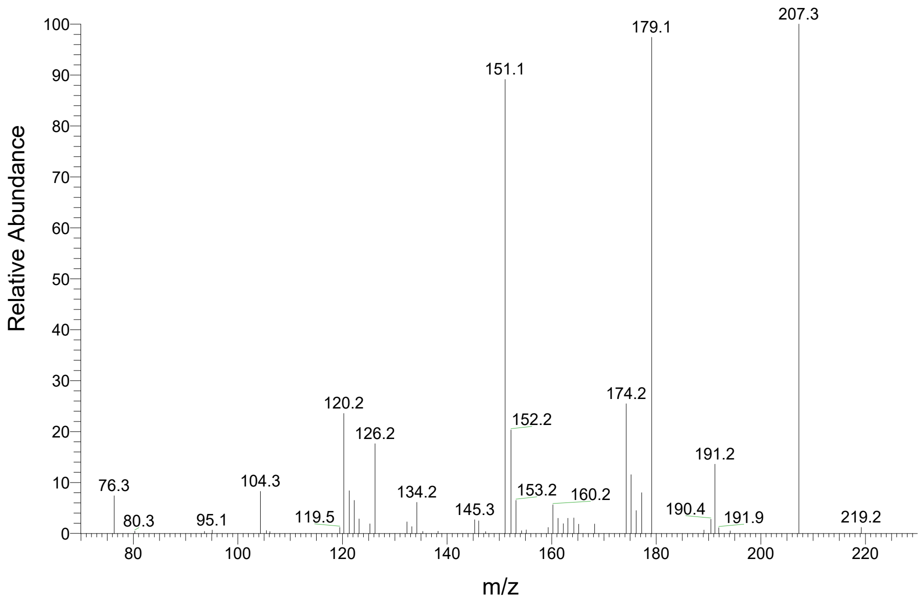

RT (min) - 10.98, $[M-H]^-$ = 237.06

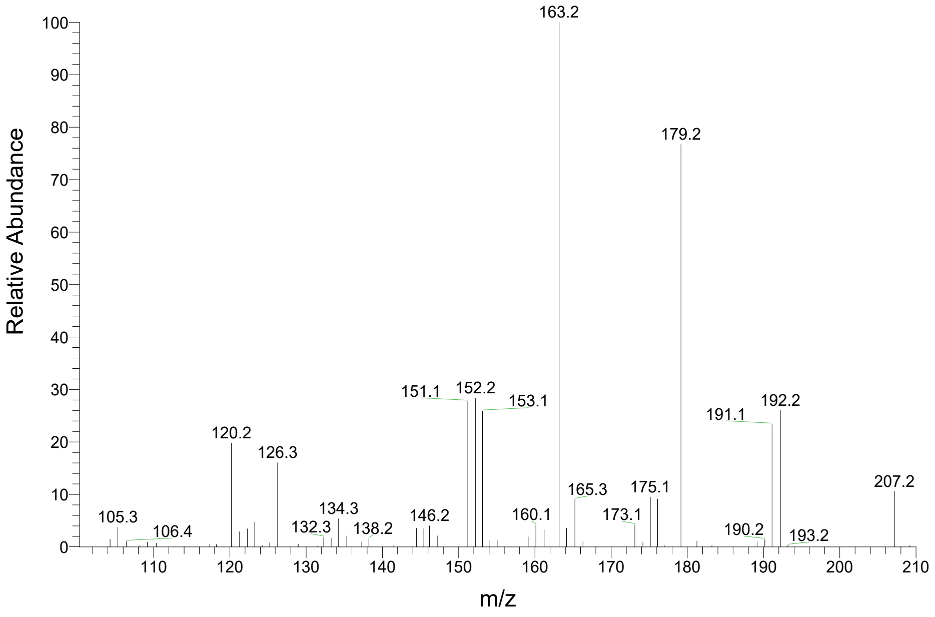


RT (min) - 15.53, $[M-H]^-$ = 251.08

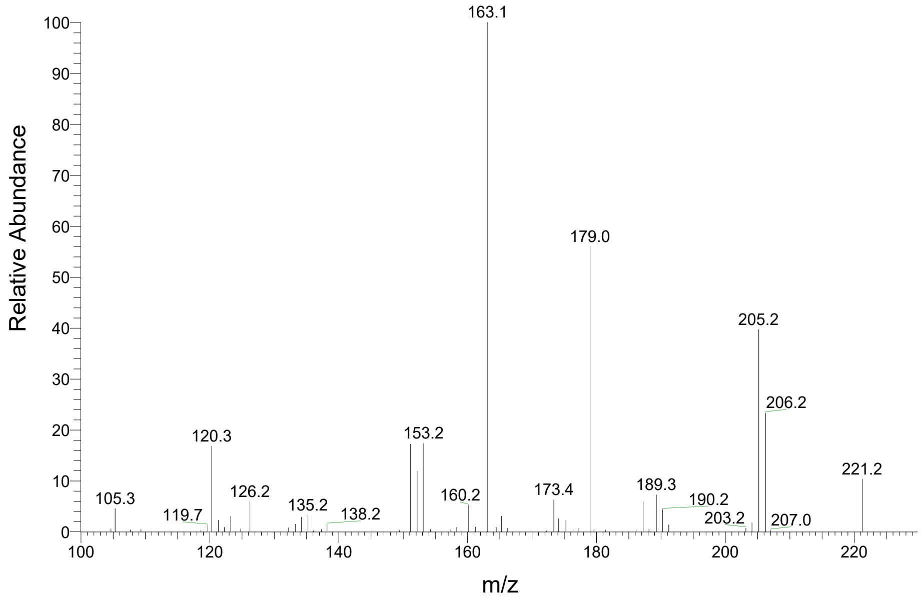


RT (min) - 17.91, $[M-H]^-$ = 299.08

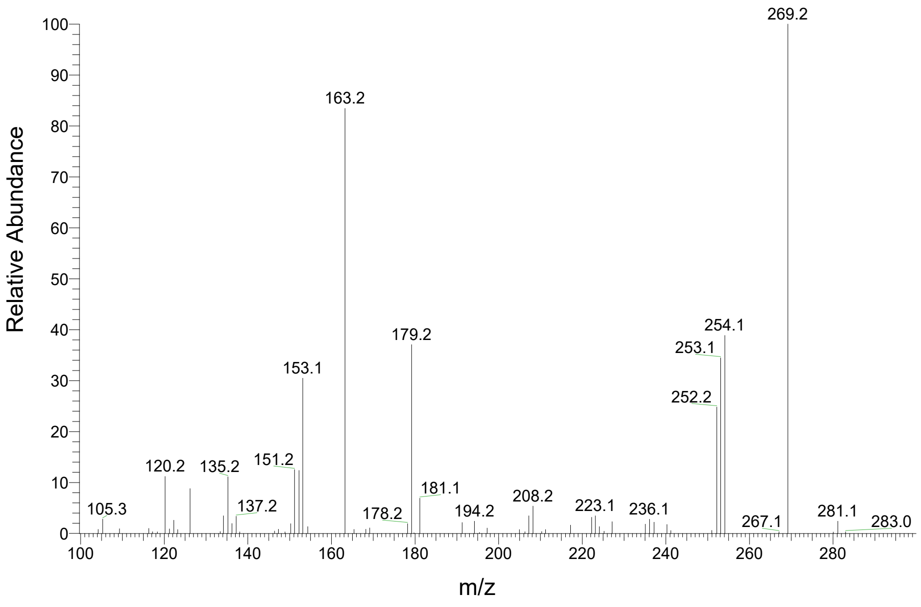


RT (min) - 18.36, $[M-H]^-$ = 263.08

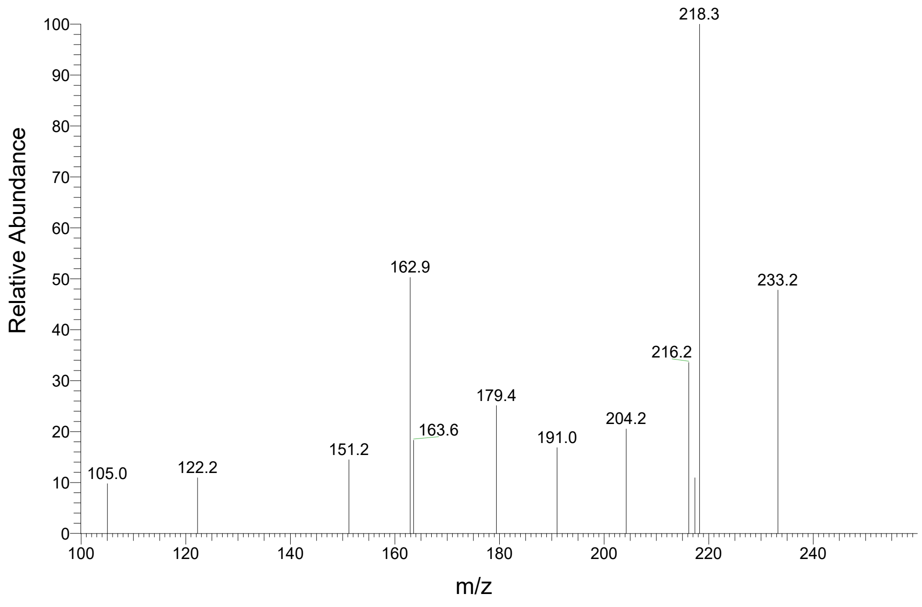

RT (min) - 19.04, $[M-H]^-$ = 265.09

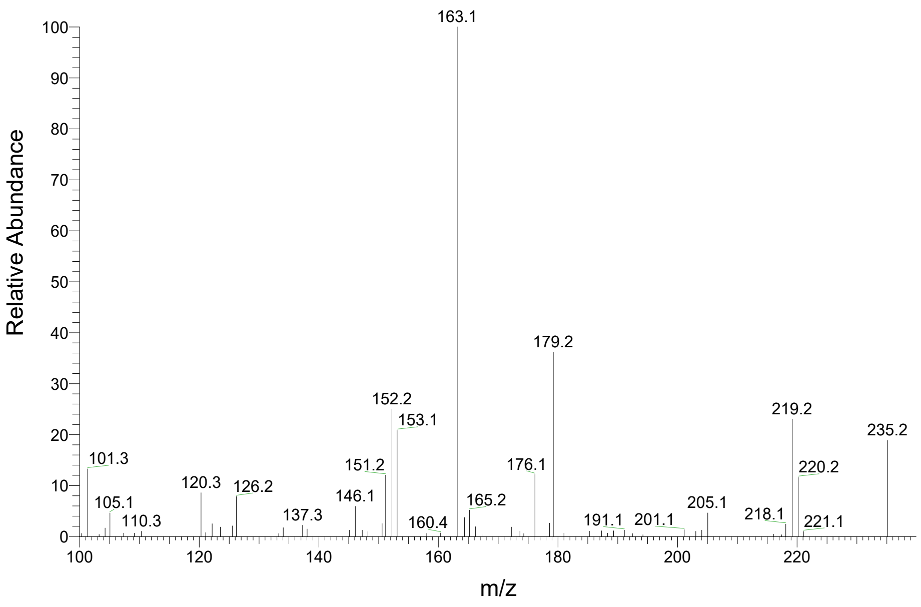


RT (min) - 20.31, $[M-H]^-$ = 277.09

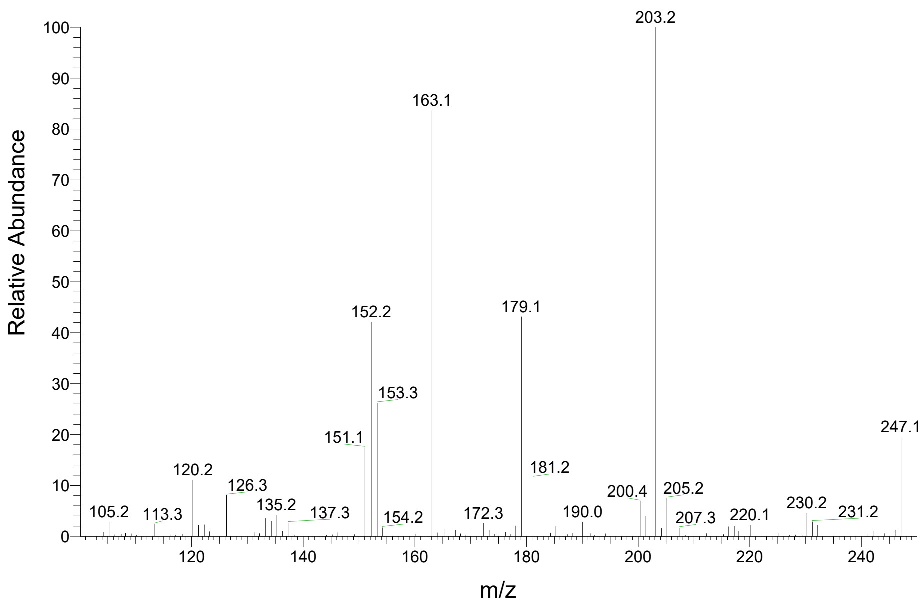


RT (min) - 22.07, $[M-H]^-$ = 277.09

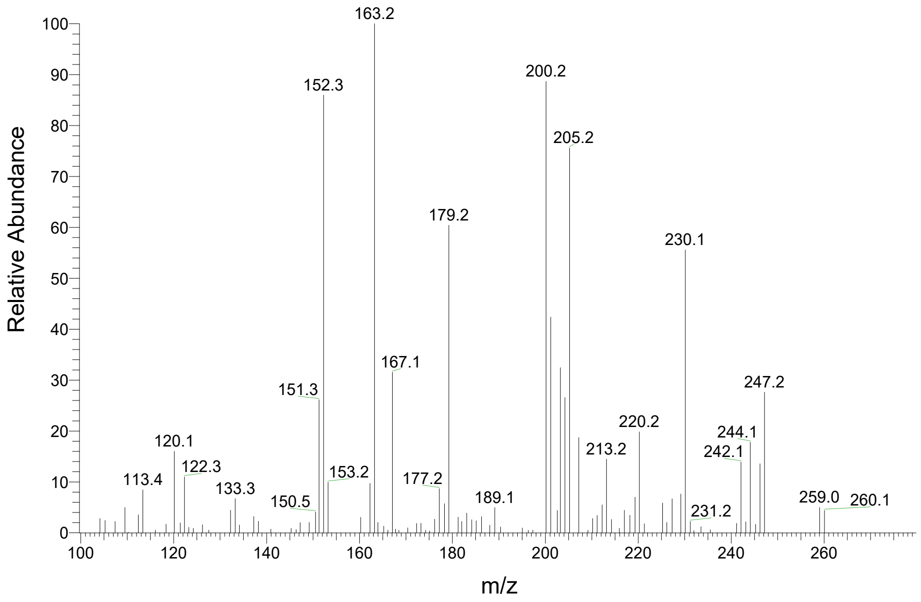


RT (min) - 23.26, $[M-H]^-$ = 279.11

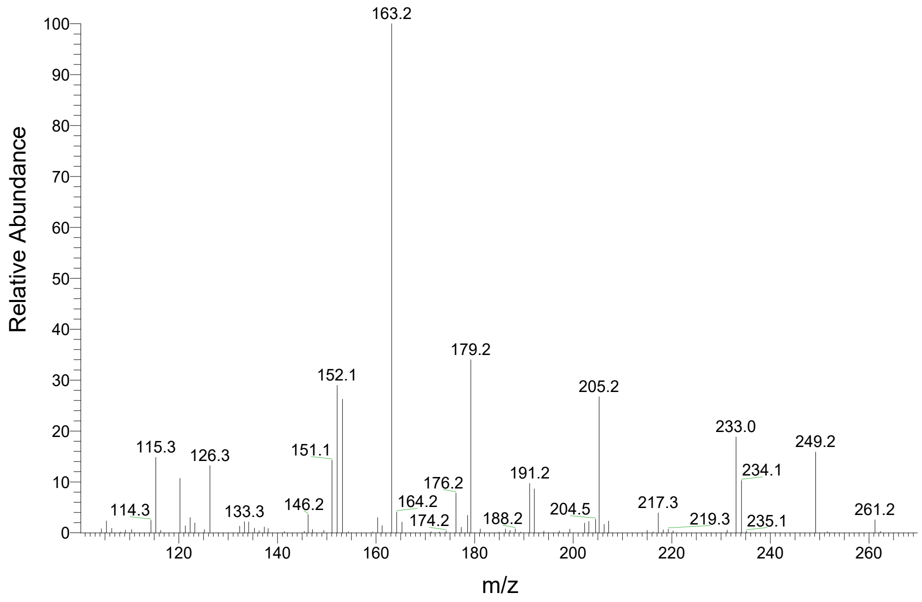

RT (min) - 35.15, [M-H]$^-$ = 321.16

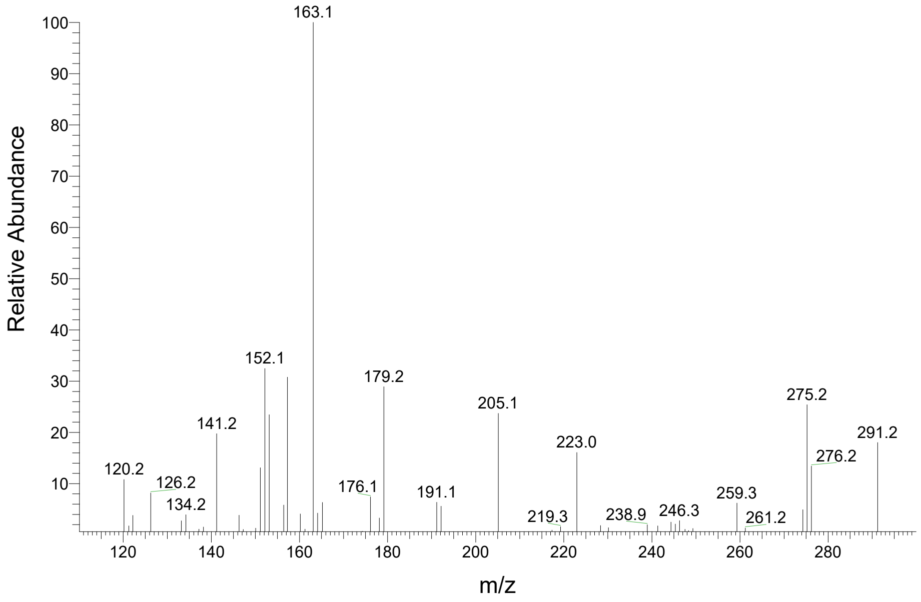
