## Supplementary Fig. S2 for "Oxylipin carbonyl composition in the chloroplast compartments"

Standard

DNP-formaldehyde ($C_7H_6N_4O_4$), $[M-H]^-$ = 209.03

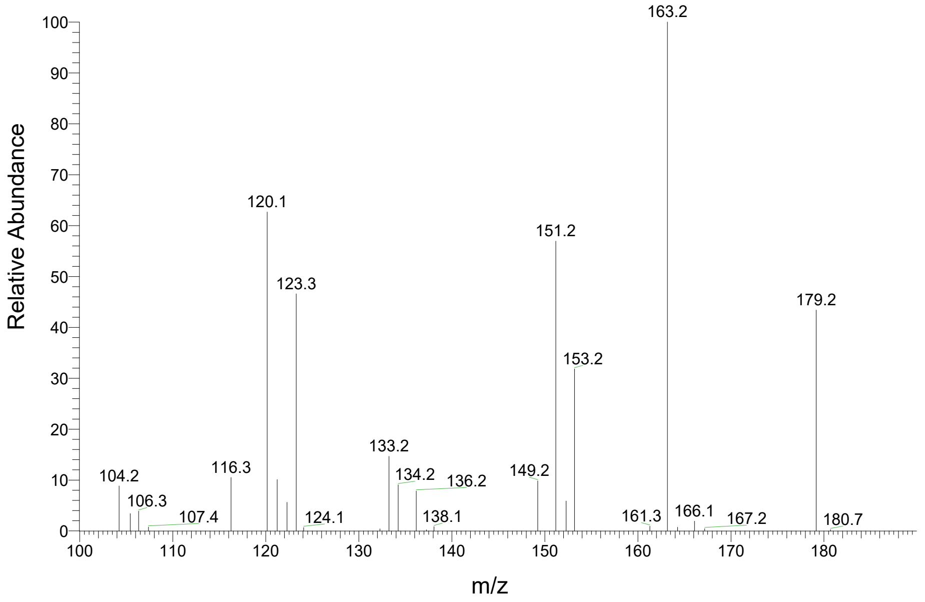


Standard

DNP-acetaldehyde ($C_8H_7N_4O_4$), $[M-H]^-$ = 223.05

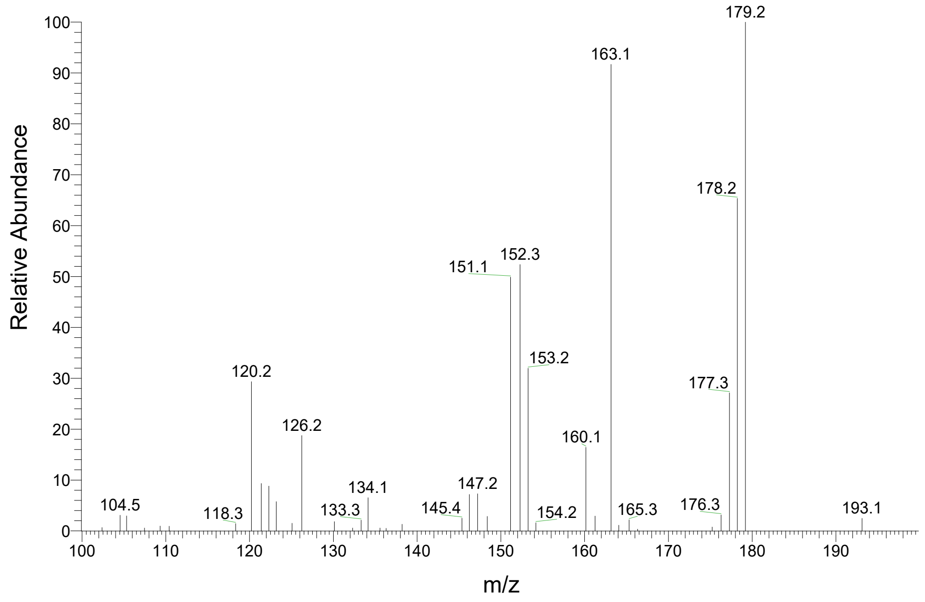


Standard

DNP-4-hydroxy-(E)-2-hexenal ($C_{12}H_{13}N_4O_5$), $[M-H]^-$ = 293.09

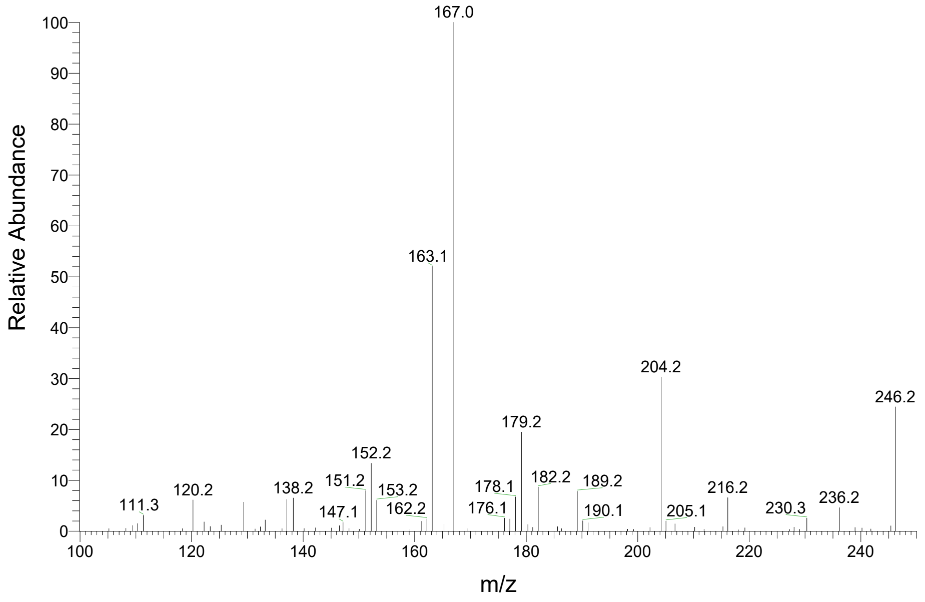


Standard

DNP-acetone ($C_9H_9N_4O_4$), $[M-H]^-$ = 237.06

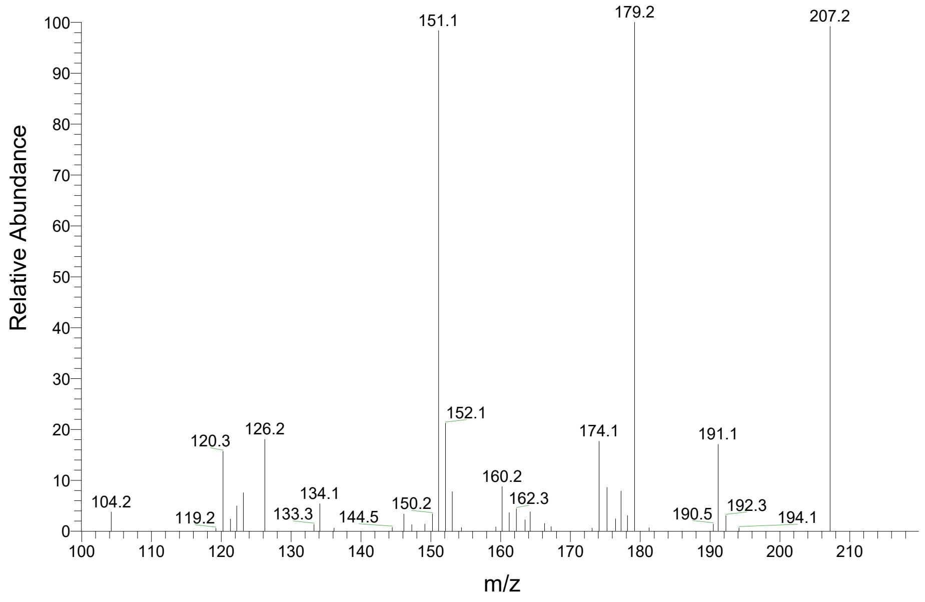

Standard

DNP-acrolein ($C_9H_7N_4O_4$), $[M-H]^-$ = 235.05

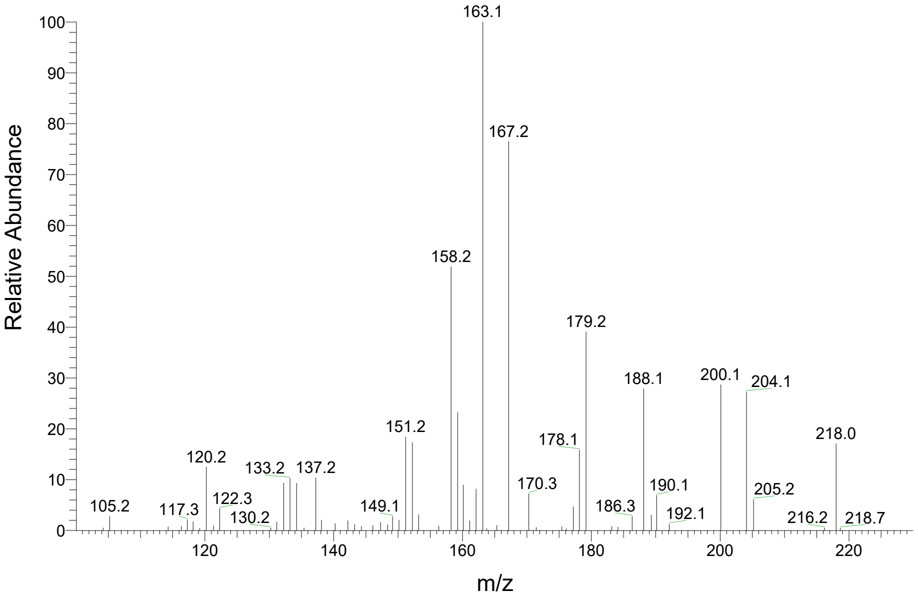


Standard

DNP-propionaldehyde ($C_9H_9N_4O_4$), $[M-H]^-$ = 237.06

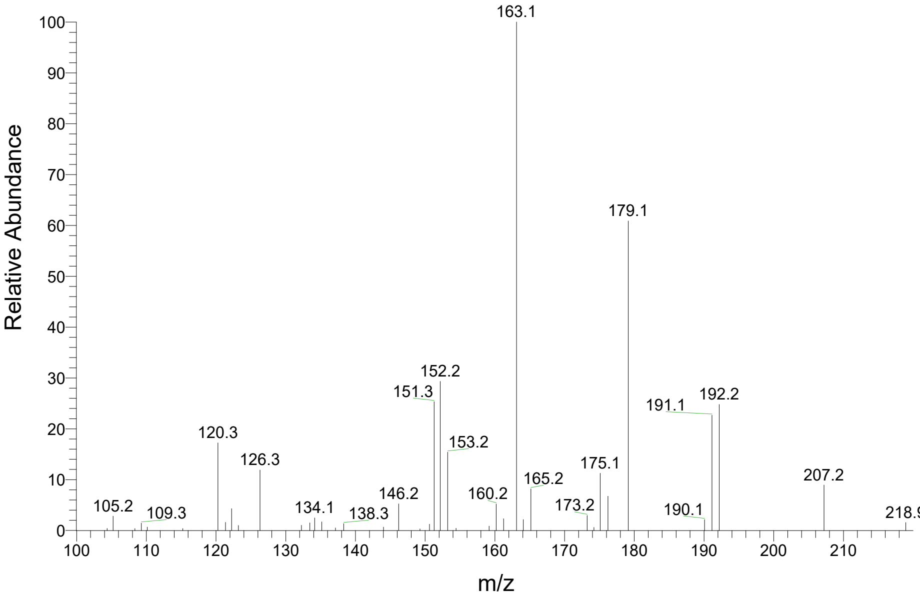


Standard

DNP-butyraldehyde ($C_{10}H_{11}N_4O_4$), $[M-H]^-$ = 251.08

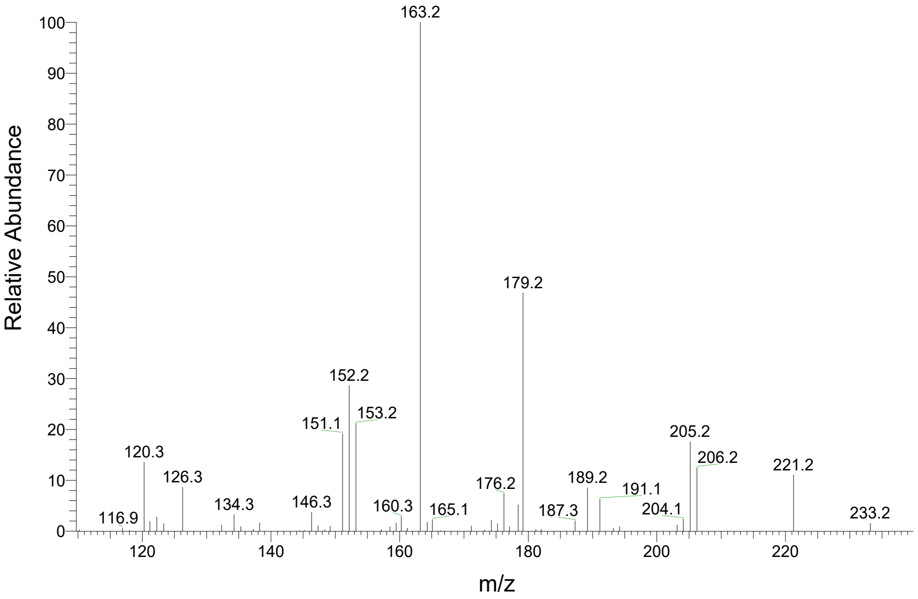


Standard

DNP-phenylacetaldehyde ($C_{14}H_{11}N_4O_4$), $[M-H]^-$ = 299.08

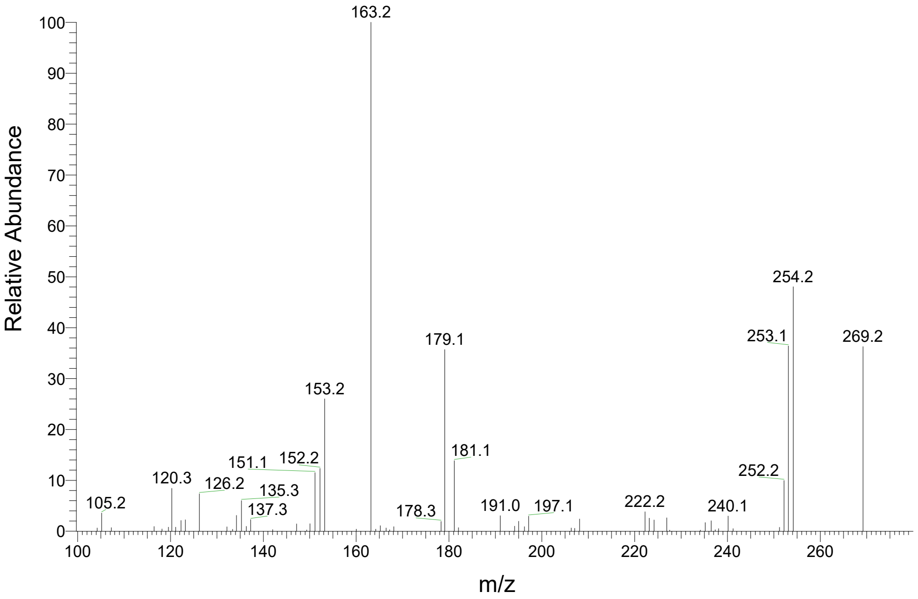

Standard

DNP-(*E*)-2-pentenal ($C_{11}H_{11}N_4O_4$), $[M\text{-}H]^-$ = 263.08

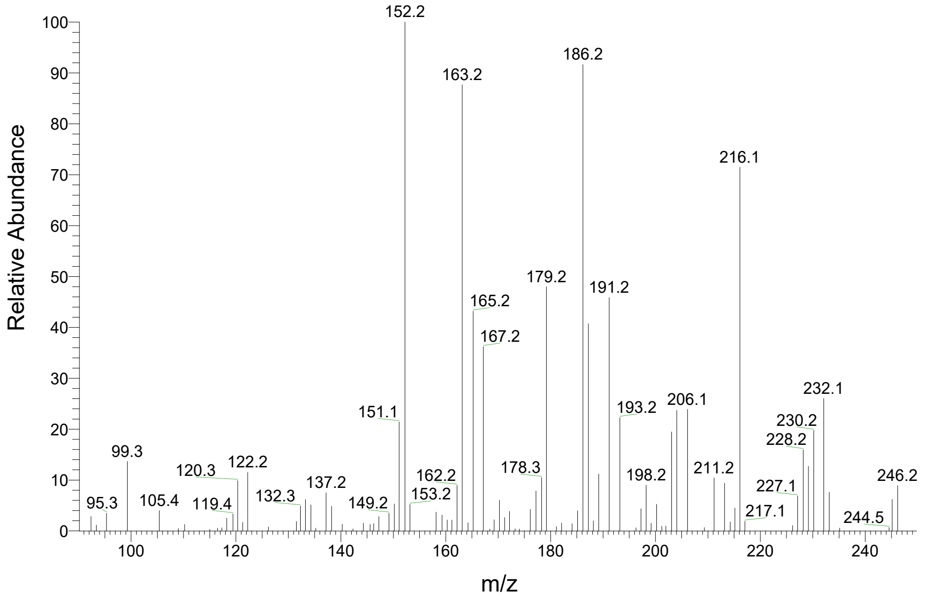


Standard

DNP-4-hydroxy-(E)-2-nonenal ($C_{15}H_{19}N_4O_5$), $[M\text{-}H]^-$ = 335.14

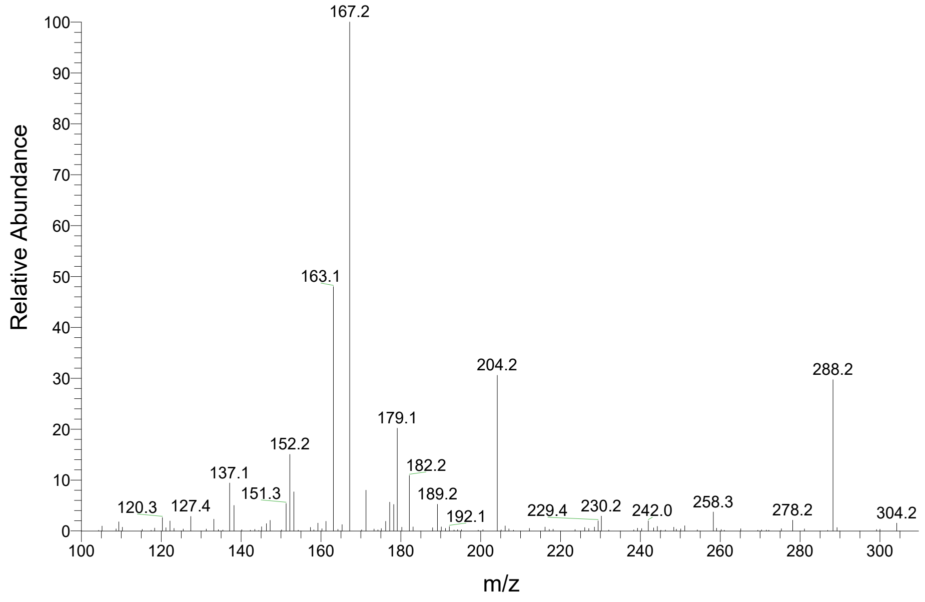


Standard

DNP-n-pentanal ($C_{11}H_{13}N_4O_4$), $[M\text{-}H]^-$ = 265.09

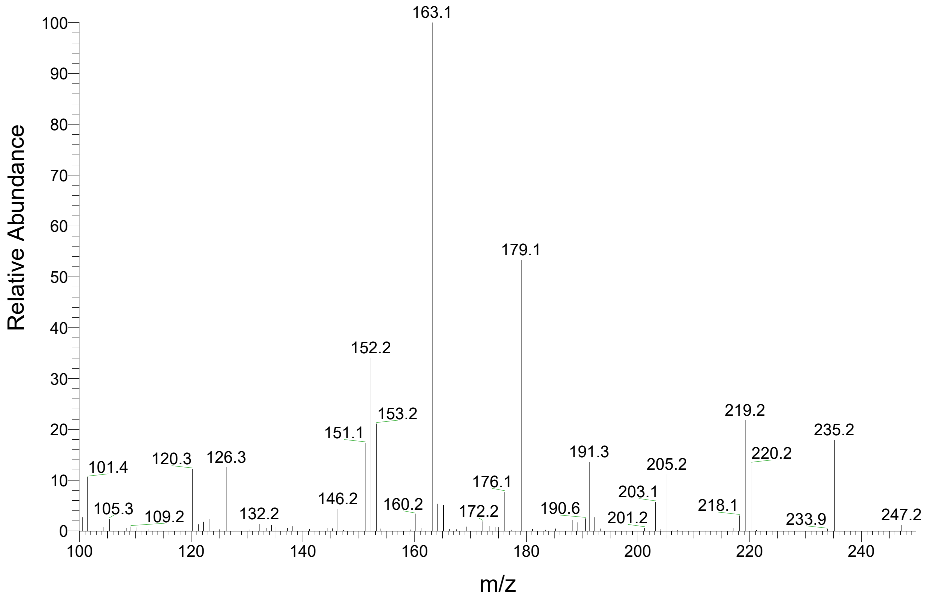


Standard

DNP-(*Z*)-3-hexenal ($C_{12}H_{13}N_4O_4$), $[M\text{-}H]^-$ = 277.09

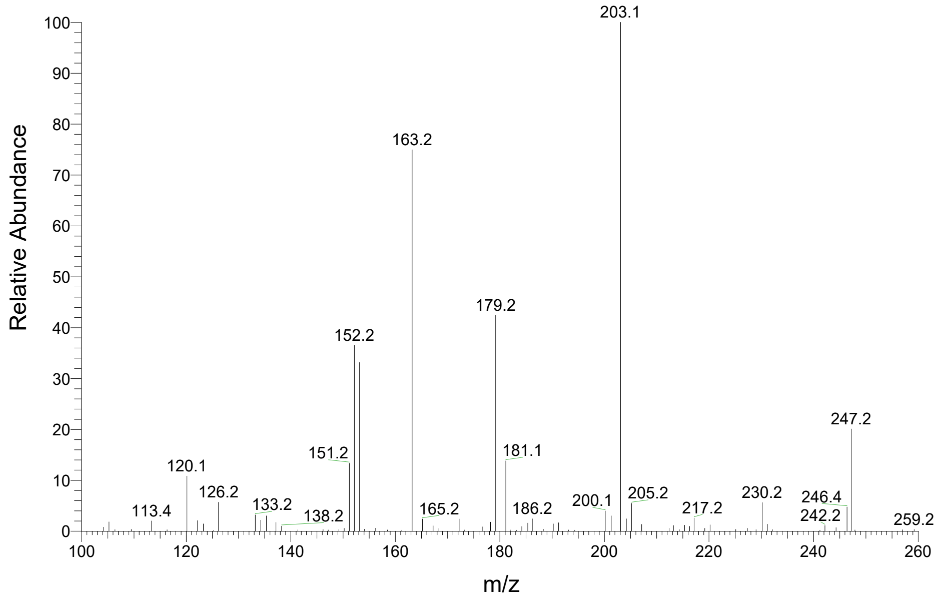

Standard

DNP-(*E*)-2-hexenal ($C_{12}H_{13}N_4O_4$), $[M-H]^-$ = 277.09

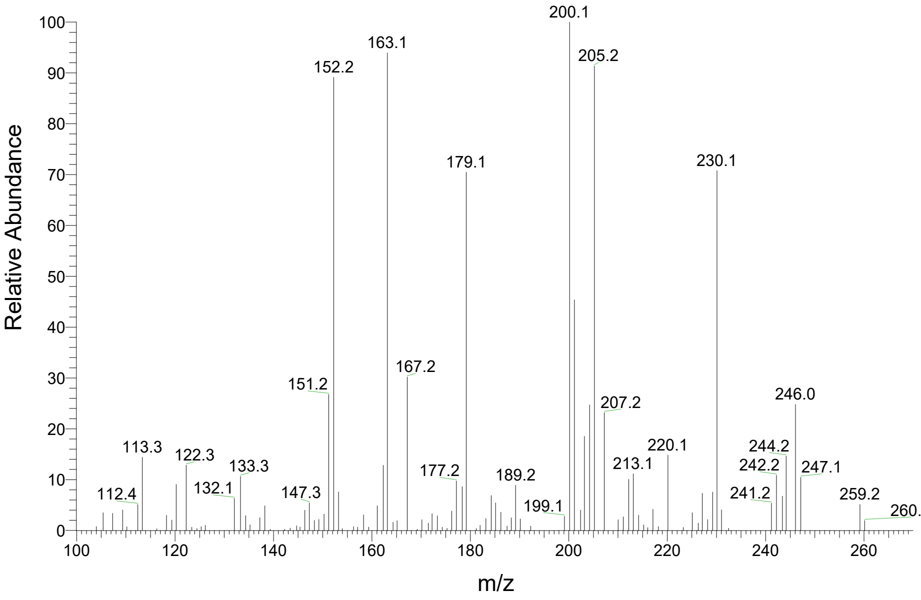


Standard

DNP-n-hexanal ($C_{12}H_{15}N_4O_4$), $[M-H]^-$ = 279.11

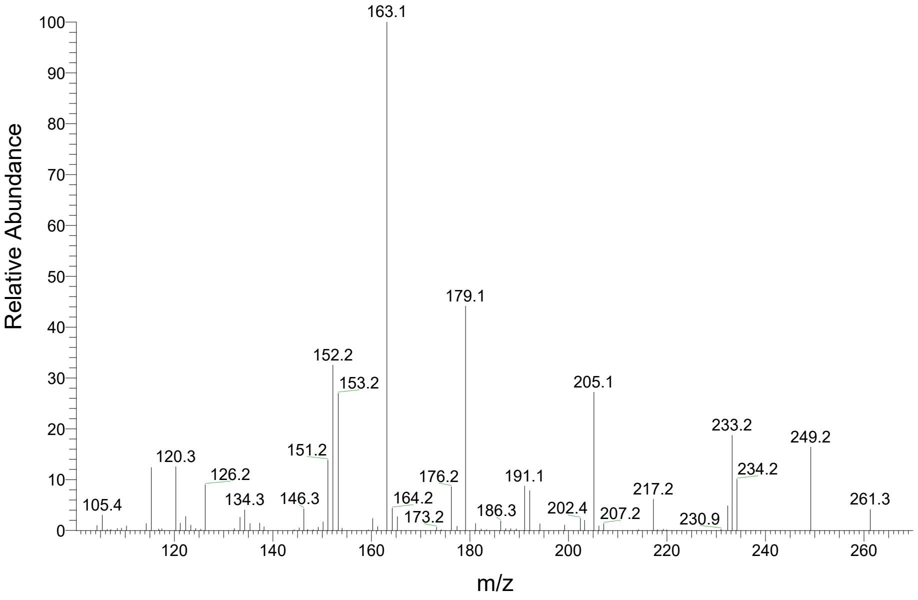


Standard

DNP-(E)-2-nonenal ($C_{15}H_{19}N_4O_4$), $[M-H]^-$ = 319.14

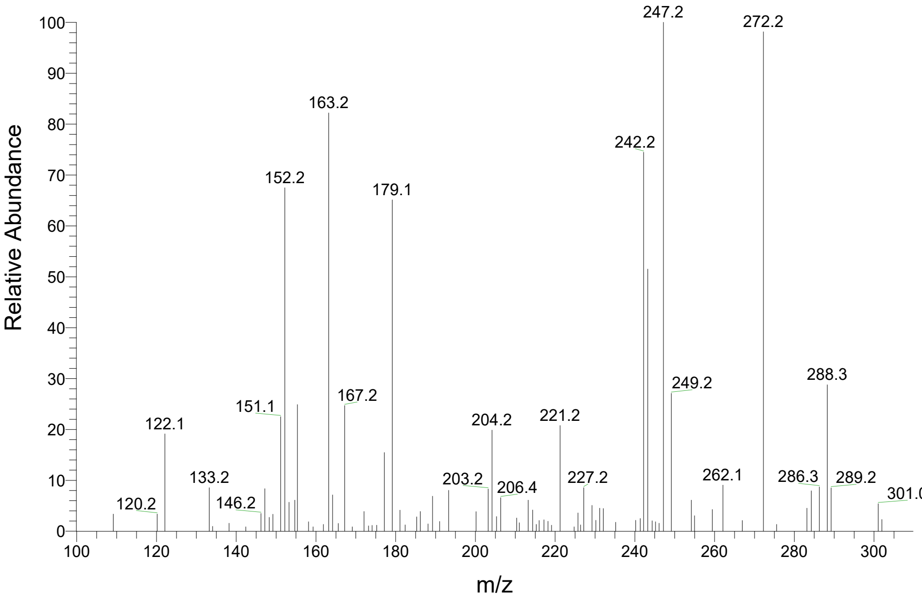


Standard

DNP-n-nonanal ($C_{15}H_{21}N_4O_4$), $[M-H]^-$ = 321.16

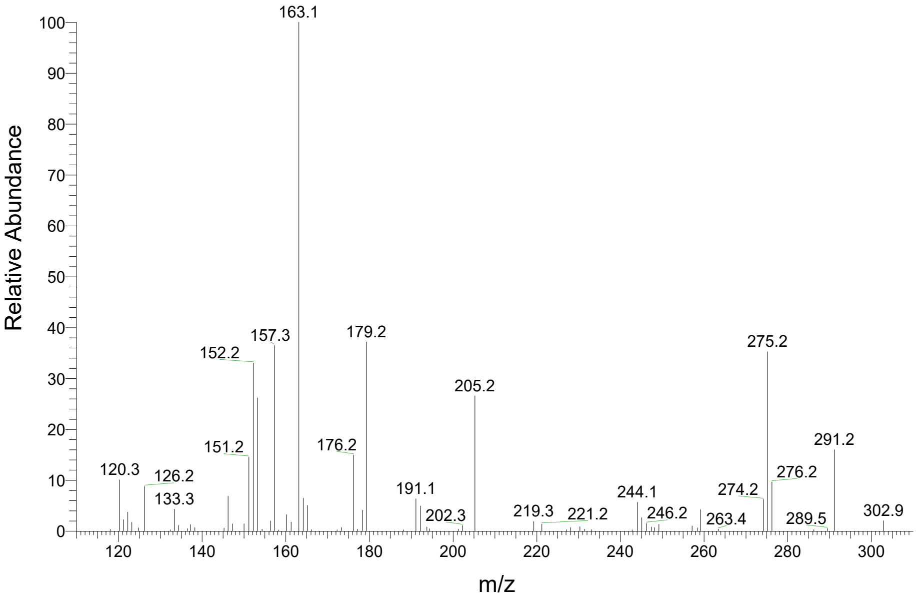
