## Supplementary Fig. S3 for "Oxylipin carbonyl composition in the chloroplast compartments"

DNP-unidentified #1 ($C_{11}H_{13}N_4O_5$), $[M-H]^-$ = 281.08

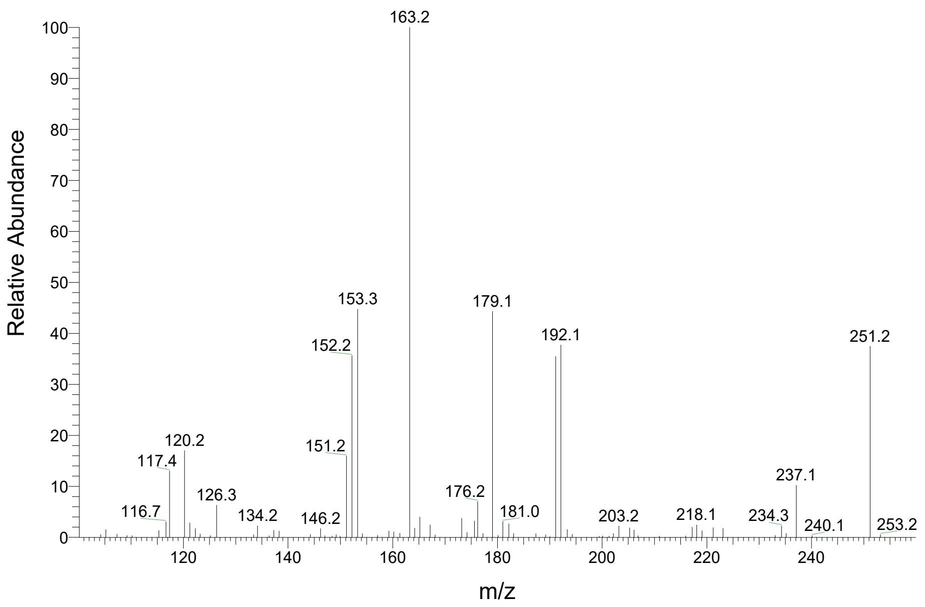


DNP-unidentified #2 ($C_{17}H_{21}N_4O_6$), $[M-H]^-$ = 377.15

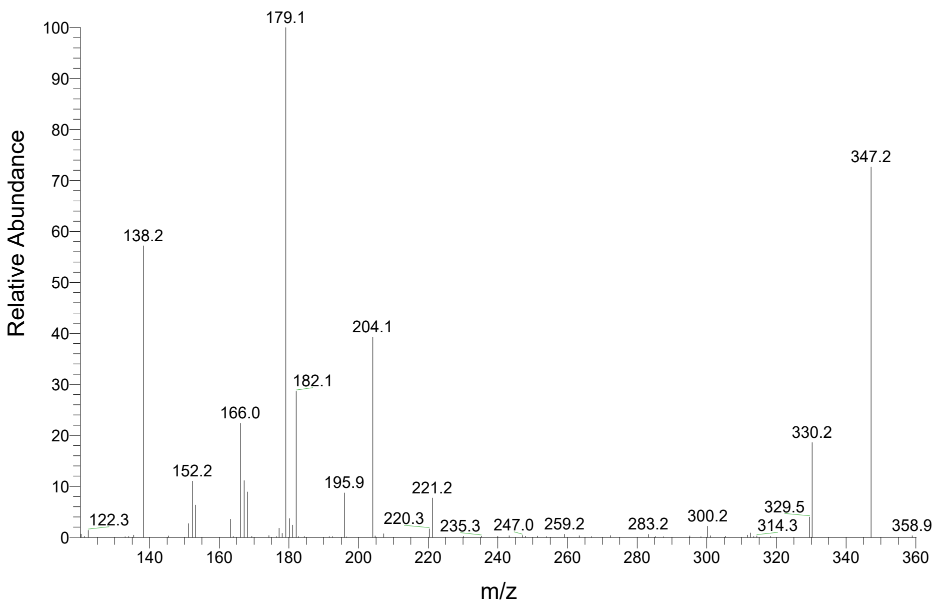


DNP-unidentified #3 ($C_{12}H_{11}N_4O_5$), $[M-H]^-$ = 291.07

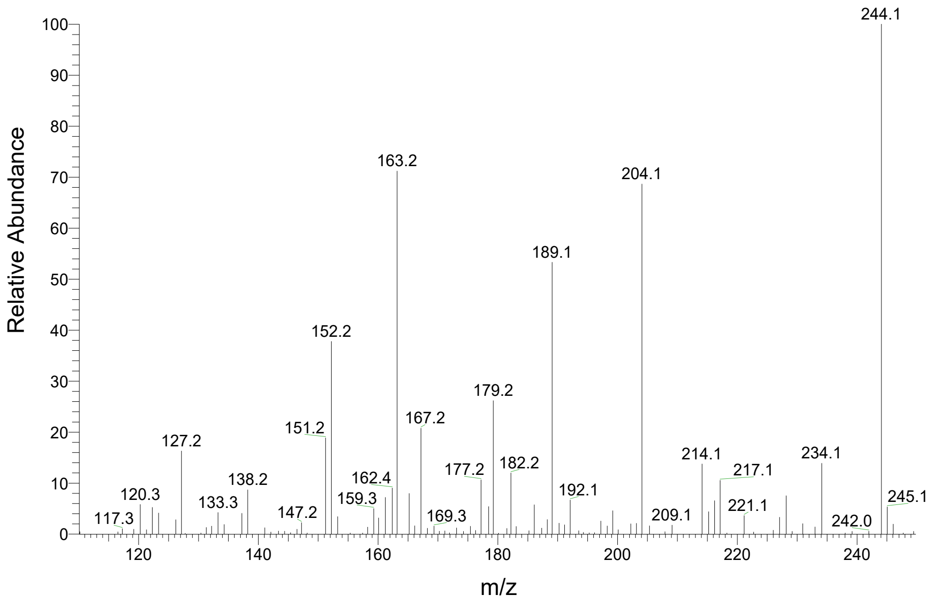


DNP-unidentified #4 ($C_{12}H_{15}N_4O_5$), $[M-H]^-$ = 295.1

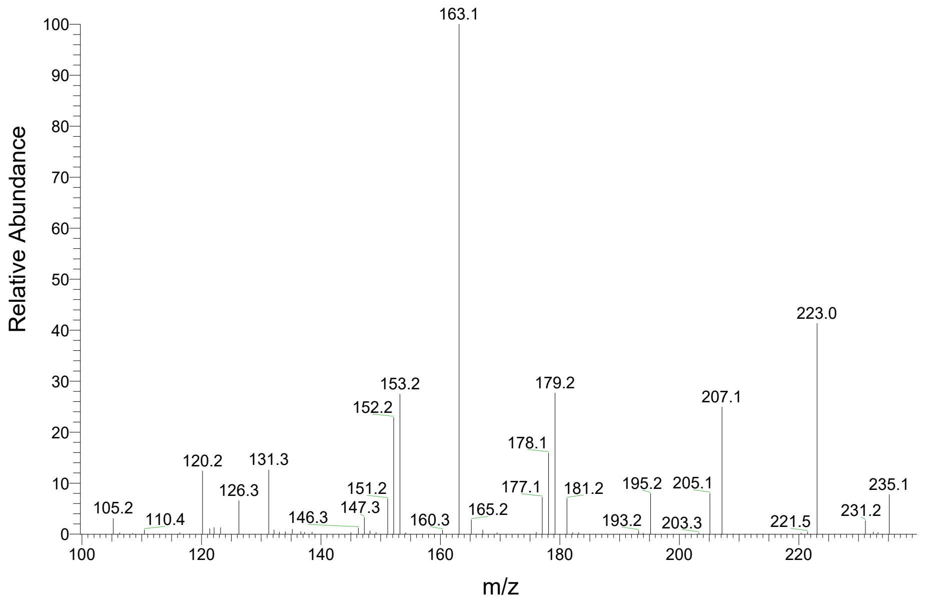

DNP-unidentified #5 ($C_{14}H_{11}N_4O_5$), $[M-H]^-$ = 315.07

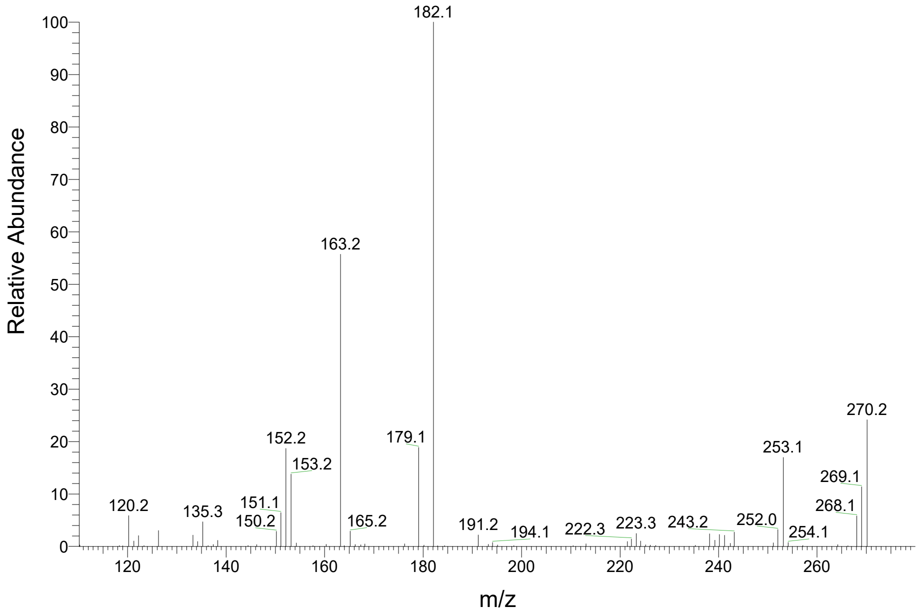


DNP-unidentified #6 ($C_{13}H_9N_4O_5$), $[M-H]^-$ = 301.06

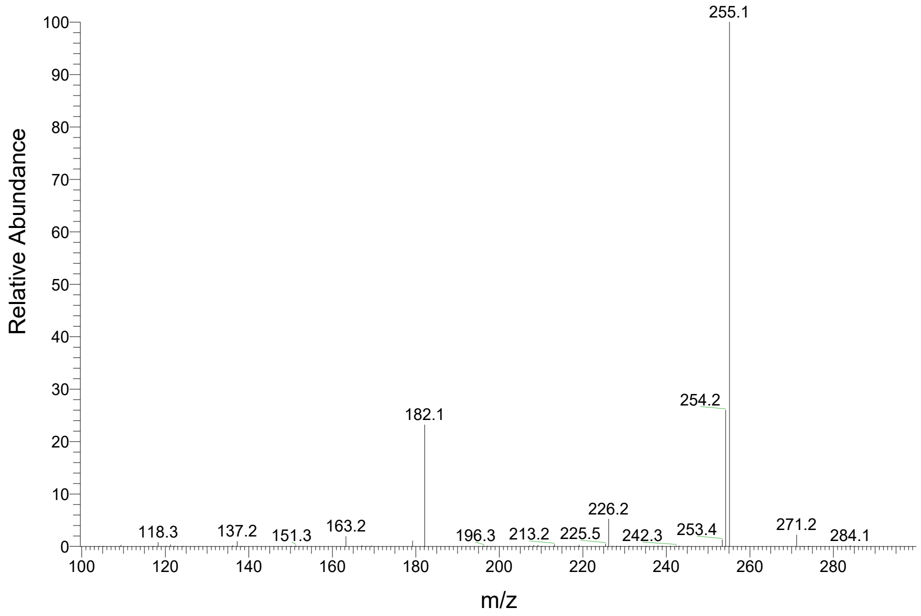


DNP-unidentified #7 ($C_{12}H_{13}N_4O_6$), $[M-H]^-$ = 309.08

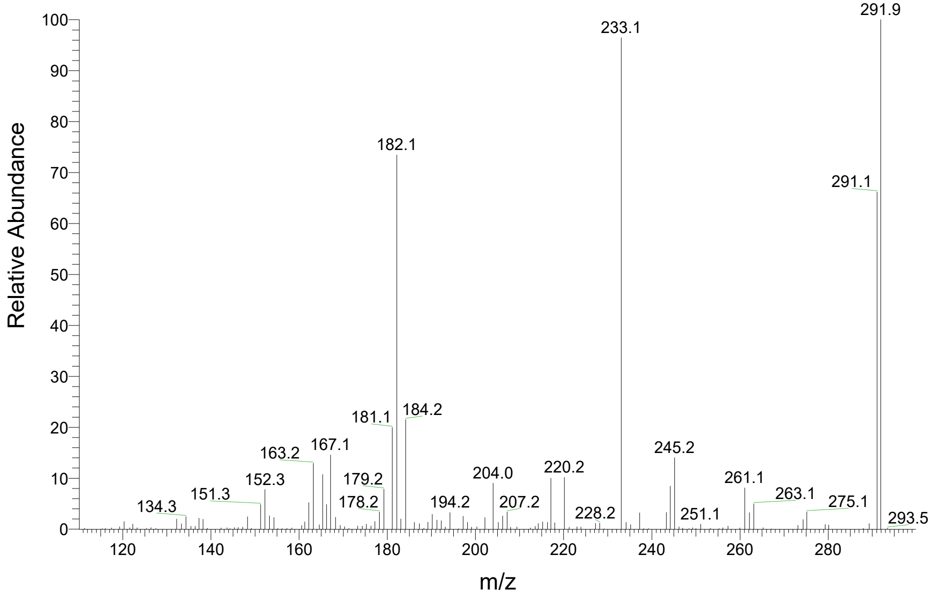


DNP-unidentified #8 ($C_{19}H_{23}N_4O_6$), $[M-H]^-$ = 403.16

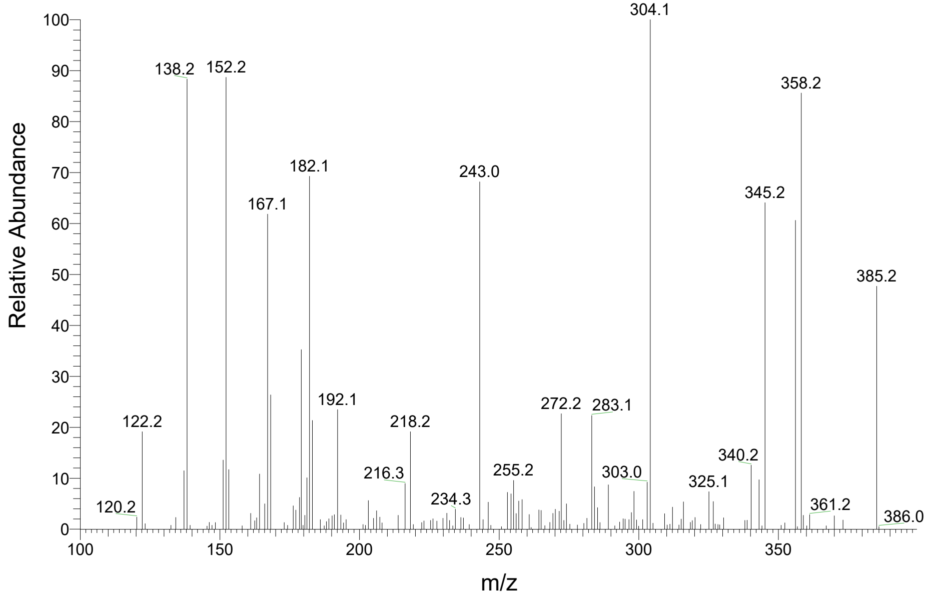

DNP-unidentified #9 ($C_{18}H_{23}N_4O_6$), $[M-H]^-$ = 391.16

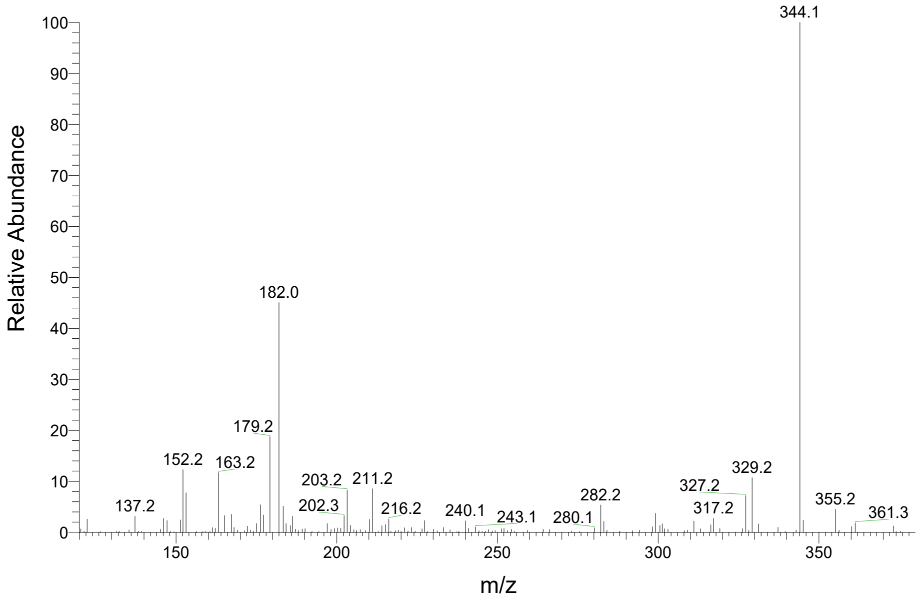


DNP-unidentified #10 ($C_{16}H_{12}N_5O_4$), $[M-H]^-$ = 338.09

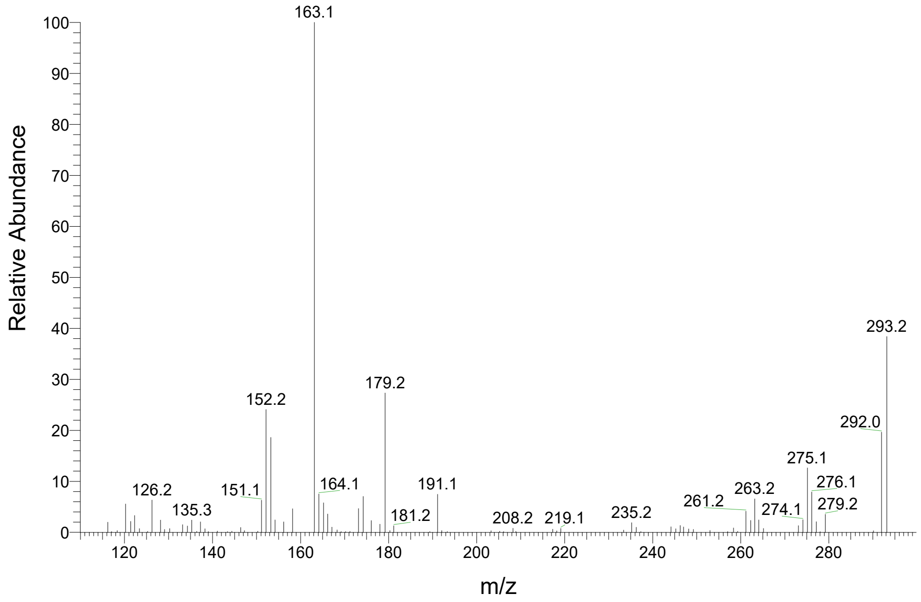


DNP-unidentified #11 ($C_{19}H_{23}N_4O_6$), $[M-H]^-$ = 403.16

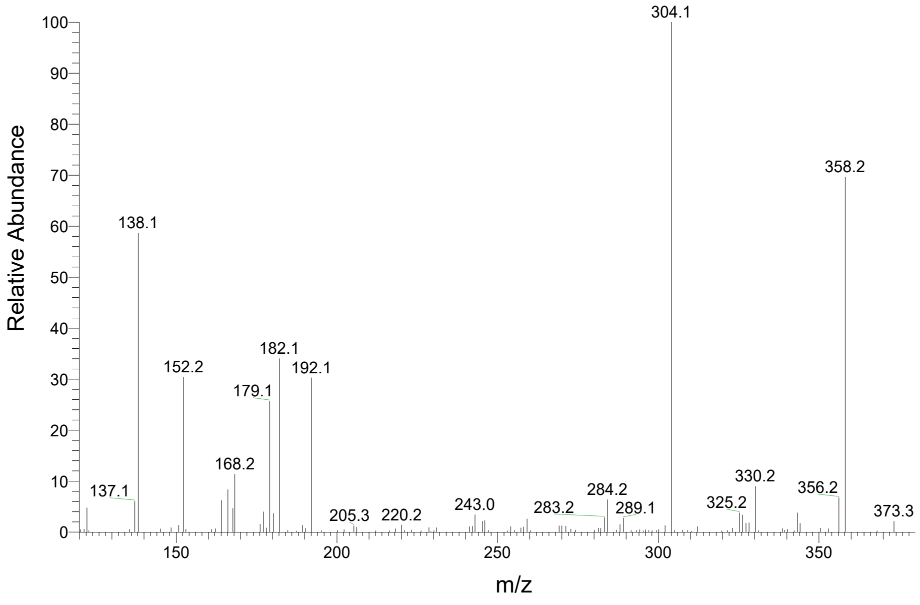


DNP-unidentified #12 ($C_{16}H_{23}N_4O_4$), $[M-H]^-$ = 335.17

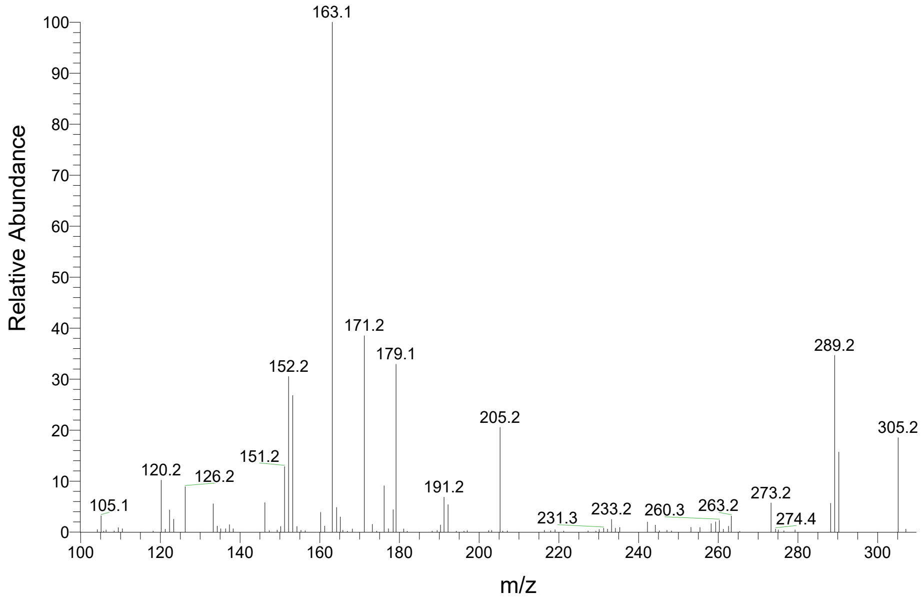
